# Consistent Consideration of RNA Structural Alignments Improves Prediction Accuracy of RNA Secondary Structures

**DOI:** 10.1101/2020.07.14.199893

**Authors:** Masaki Tagashira

## Abstract

The probabilistic consideration of the global pairwise sequence alignment of two RNAs tied with their global single secondary structures, or global pairwise structural alignment, is known to predict more accurately global single secondary structures of unaligned homologs by discriminating between conserved local single secondary structures and those not conserved. However, conducting rigorously this consideration is computationally impractical and thus has been done to decompose global pairwise structural alignments into their independent components, i.e. global pairwise sequence alignments and single secondary structures, by conventional methods. ConsHomfold and ConsAlifold, which predict the global single and consensus secondary structures of unaligned and aligned homologs considering consistently preferable (or sparse) global pairwise structural alignments on probability respectively, were developed and implemented in this study. These methods demonstrate the best trade-off of prediction accuracy while exhibiting comparable running time compared to conventional methods. ConsHomfold and ConsAlifold optionally report novel types of loop accessibility, which are useful for the analysis of sequences and secondary structures. These accessibilities are average on sparse global pairwise structural alignment and can be computed to extend the novel inside-outside algorithm proposed in this study that computes pair alignment probabilities on this alignment.

## INTRODUCTION

Investigating the (global secondary) structures of potentially functional *ncRNA*s is an important key to discover functional ncRNAs and uncover their functional details, because single structures are often conserved among homologs even in the case where the sequences of these homologs are nonconserved. (1, 2, 3) Methods that predict a structure from only a sequence (or *mono-folding*s), such as RNAfold (4), Mfold (5), Pfold (6), Sfold (7), CONTRAfold (8), and CentroidFold (9), have aided experimental single structure probing methods, such as Cryo-EM (10), icSHAPE (11), SHAPE-MaP (12), PARS (13), and PARIS (14), to provide input model single structures and predict single structures constrained by single structure probing data (15, 16, 17).

Mono-foldings suffer from the limited accuracy of predicted single structures, because there is no room for exploiting information other than single sequences. (Figure 1c) In order to predict more accurate structures, other sequences that share homology with a sequence are used. Predicting the (global) sequence alignment of RNAs tied with their single structures, or (global) *structural alignment* (18), is one of the most effective ways to predict more precise structures of homologs, which are informed by (base-)pairing substitutions (1, 19), though many methods that predict structural alignments report *consensus structure*s, rather than single structures. Consensus structures are a set of column pairs on sequence alignment that contain pairings conserved among homologs. The serious problem of predicting rigorous structural alignments is the impractical running time and memory usage of this prediction, even if this prediction is pairwise, which are bounded by *O*(*L*^3*R*^) and *O*(*L*^2*R*^), respectively, where *L* and *R* are the maximum length and the number of sequences, respectively. (18) This problem is solved by utilizing the progressive alignment (20) and *sparsification*, which filters out candidates that do not satisfy certain thresholds from consideration to set the scores of these candidates to −∞, by many popular methods that predict structural alignments, such as PMcomp (21), Foldalign (22, 23), Murlet (24), MXSCARNA (25), LocARNA (26), RAF (27), DAFS (28), and SPARSE (29).

**Figure 1.**
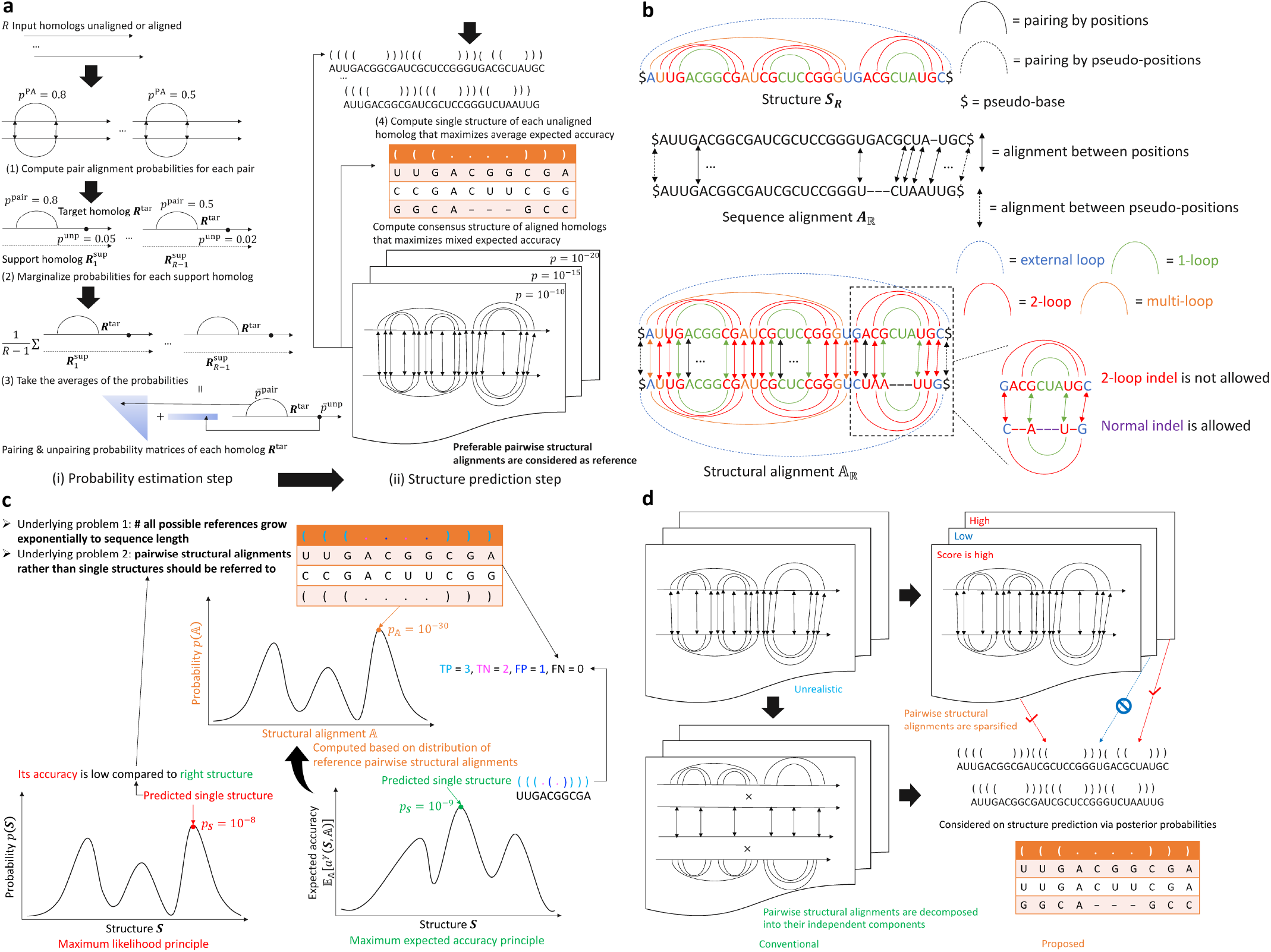
**(a) The proposed workflow of ConsHomfold and ConsAlifold.** ConsHomfold predicts single structures from its input unaligned homologs. ConsAlifold predicts a consensus structure from its input aligned homologs. **The methods do not produce structural alignments.** The methods consist of two steps to (i) estimate probabilities and (ii) predict structures after this estimation. **For ConsAlifold, the estimation step ignores the gaps in the alignment.** (1) First, pair alignment probabilities on sparse pairwise structural alignment for each pair of input homologs are computed. (2) Then, the probabilities are marginalized to be pairing and unpairing probabilities of each homolog. (3) As the final process of the estimation step, average pairing and unpairing probabilities between each homolog and the remaining homologs are gained. These probabilities are utilized to incorporate more than one support homolog into the subsequent predictions of structures. (4) Finally, ConsHomfold predicts the single structure of each input homolog and ConsAlifold predicts the consensus structure of the input alignment. **(b) Examples of a structure** *S_R_*, **a sequence alignment** 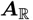, **and a structural alignment** 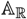. A structure ***S_R_*** and an alignment 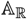 are color-coded based on types of loop. **(c) The underlying problems on conventional mono-foldings and ali-foldings.** The maximum likelihood principle, which is equivalent to the free energy minimization on structure predictions (e.g. RNAfold and RNAalifold), exerts less prediction accuracy in general compared to the maximum expected accuracy principle, which is based on Bayes’ theorem (e.g. CONTRAfold, CentroidFold, CentroidHomfold, TurboFold, PETfold, CentroidAlifold, ConsHomfold, and ConsAlifold), because the most probable references have their probabilities, which decay exponentially to sequence length. CONTRAfold, CentroidFold (conventional mono-foldings), PETfold, and CentroidAlifold (conventional ali-foldings) consider not pairwise structural alignments but single structures. **(d) Considering all possible pairwise structural alignments costs** *O*(*N*^3^*M*^3^) **long hours and** *O*(*N*^2^*M*^2^) **huge memory where** *N* **and** *M* **are the lengths of homologs.** The conventional hom-foldings, CentroidHomfold and TurboFold, decompose all pairwise structural alignments into their independent pairwise sequence alignments and single structures. **ConsHomfold and ConsAlifold take consistently sparse pairwise structural alignments into account.**

As other ways to predict more precise structures from homologs, the following methods are available:

- *hom-folding*: predicting the single structure of each unaligned homolog considering pairwise structural alignments between this homolog and the remaining unaligned homologs on the probability distributions of these alignments as CentroidHomfold (30) and TurboFold (31)
- *ali-folding*: predicting the consensus structure of aligned homologs as RNAalifold (32), PETfold (33), and CentroidAlifold (34).

The above foldings are more reasonable than structural alignment predictions, because the progressive alignment, which is not required for these foldings, is computationally complex and heavy. The conventional hom-foldings (CentroidHomfold and TurboFold) decompose pairwise structural alignments on probability into their independent components, i.e. pairwise sequence alignments and single structures. (Figure 1d) Also, the conventional ali-foldings (RNAalifold, PETfold, and CentroidAlifold) do not consider pairwise structural alignments on probability, which will improve the prediction accuracy of these ali-foldings. (Figure 1c) There is possibly room for improving the prediction accuracy of hom-foldings and ali-foldings to exploit more accurately the homology among homologs.

### Contributions of this study

A hom-folding and an ali-folding that consider consistently likely (or sparse) pairwise structural alignments on probability were proposed in this study as ConsHomfold and ConsAlifold (Figure 1a), respectively. (Figure 1d) These foldings are of the maximum expected accuracy principle whose prediction is in general more accurate than that of the maximum likelihood principle, because the number of all possible references grow exponentially to sequence length and thus results in the extremely high dimension of predictive space (35, 36). (Figure 1c) ConsHomfold and ConsAlifold compute *pair alignment probabilities* (or *pair match probabilities*) on sparse pairwise structural alignment that were proposed including the computation of them in LocARNA-P (37). The scores to calculate these probabilities in these foldings are based on *Turner’s (nearest neighbor physics) model*, which scores single structures in terms of the free energy of their *loop*s (38), and are expected to be more suitable to score structural alignments than those that are used in LocARNA-P. An algorithm to compute the probabilities was developed in this study, though this algorithm shares the framework of inside-outside algorithm with LocARNA-P. The novel algorithm requires (quadratic) less computational complexities than those quartic of LocARNA-P to impose more hard sparsity.

Also, novel types of the *loop accessibility* (= *structural profile* = *structural context*) proposed in CapR (39), which is useful for the analysis of RNAs and their structures, and an algorithm to compute the novel accessibilities were proposed in this study to extend that of the novel alignment probabilities. These novel accessibilities are based on sparse pairwise structural alignment, whereas those in CapR are based on local single structure. ConsHomfold and ConsAlifold output optionally the novel accessibilities.

## MATERIALS AND METHODS

### Pairwise structural alignment

Let ***S***_***R***_, 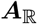, and 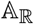 be the structure of the sequence ***R***, the sequence alignment between the pair of sequences ℝ, and the structural alignment between the pair ℝ, respectively. An alignment 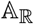 is composed of the alignment 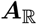 ***S***_***R***_ and ***S***_***R′***_, i.e. 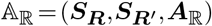 where *ℝ*=(*R*,*R*′). (Figure 1b) Assume *collinear* pariwise sequence alignments (40) and single structures without any *pseudoknot*s (41) to avoid larger computational complexities.

A position *u* is said to be *accessible from* (or *closed by*) pairing positions *i* and *j* if *i*<*u*<*j* and the position pair (*i,j*) is the closest to the position *u* of all pairing position pairs. A position set about pairing positions *i* and *j* 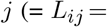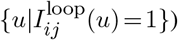 is the loop of the positions *i* and *j* where 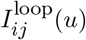 is 1 if the position *u* is accessible from the positions *i* and *j* and 0 otherwise. A loop *L*_*ij*_ is a *b-loop* if the loop *L*_*ij*_ contains *b* − 1 pairing position pairs accessible from the positions *i* and *j*. A loop *L*_*ij*_ is said to be *internal* and *external* (or *outer*) if the positions *i* and *j* are accessible from the loops of other positions and the pseudo-positions (= the virtual positions closing the both ends of the sequence ***R***), respectively. (Figure 2) (Internal) *b*-loop is divided into three classes on Turner’s model, which approximates the free energy of structure on thermodynamics (38): 1-loop (= *hairpin loop*), 2-loop (= *stacking (loop), bulge loop, and interior loop*), and (*b*>2)-loop (= *multi-loop*).

**Figure 2.**
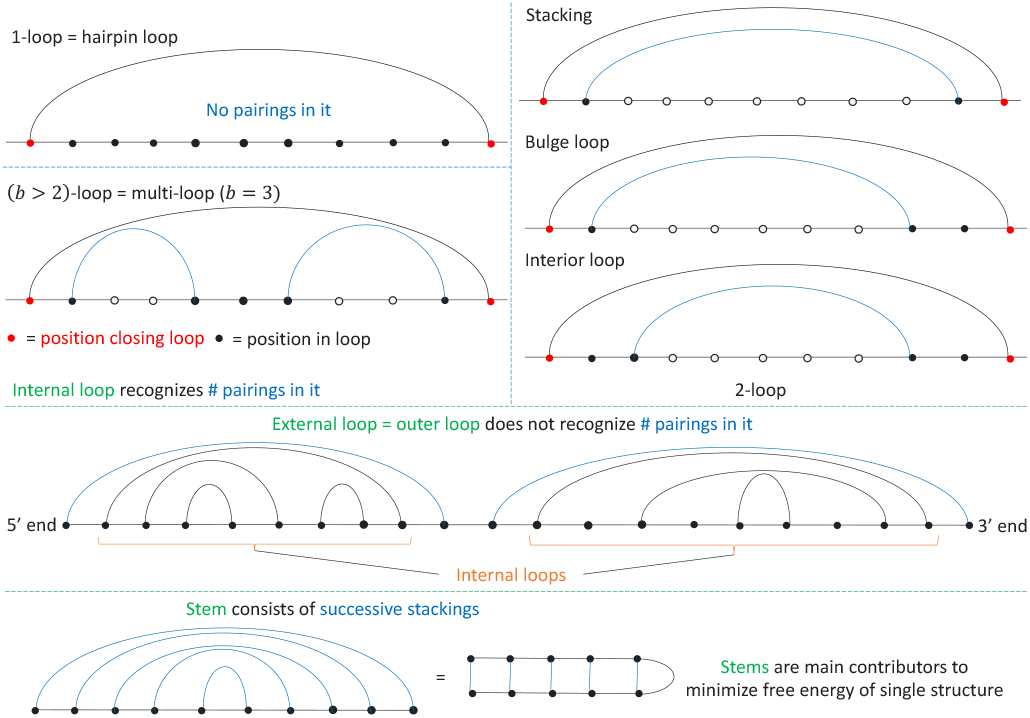
Different types of loop constitute a single structure. A stacking does not contain unpairings accessible in it. A bulge loop contains unpairings accessible in it on either of the 5’ and 3’ sides. An interior loop contains unpairings accessible in it on both of these sides.

Assume structural alignments without any *indels of 2-loop*s (Figure 1b), which are required to align *stem (loops)* (= lines of successive stackings) of different lengths keeping pairings (18, 29), to prevent scoring structural alignments from becoming further complicated though scoring these indels does not increase computational complexities (29). Two sets of position pairs are said to be *pair-aligned* if these sets are pairing and aligned. Two positions are said to be *loop-aligned* if these positions are unpaired and aligned.

### Posterior pair alignment probability matrix

Let 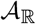 and 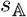 be a set of all possible alignments 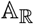 and the score of the alignment 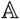, respectively. Assume that the probability of any alignment 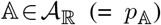 obeys a *Boltzmann (probability) distribution*, i.e. 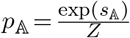 where 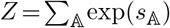. *Z* is called a *partition function*.

Let 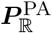 be the *pair alignment probability* matrix given the pair ℝ. Let 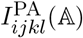 be 1 if the pairs (*i,j*) and (*k,l*) are pair-aligned in the alignment 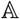 and 0 otherwise, where *i* and *j* are two positions in the sequence ***R***, *k* and *l* are two positions in the other sequence ***R***′, *i*<*j*, and *k*<*l*. The matrix 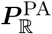 can be written by the probabilities 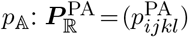 where 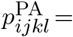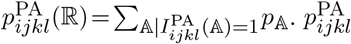 is the probability that the pairs (*i,j*) and (*k,l*) are pair-aligned.

### Composition of pairwise structural alignment score 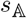

Let *e*_***S***_ be the free energy of the structure ***S***. Score 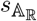 is decomposed into additional components: 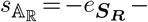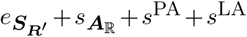. Here, 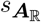, *s*^PA^, and *s*^LA^ are the score of the alignment 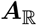, the sum score of the pair alignments in the alignment 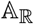, and the sum score of the loop alignments in the alignment 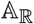, respectively.

The components 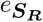 and 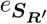 can be computed by the estimated parameters of Turner’s model. On it, energy *e*_***S***_ is decomposed into four categories of additional component: 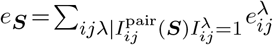 where *λ* ∈{1,2,multi,outer} and 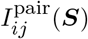 returns 1 if the positions *i* and *j* are pairing in the structure ***S***. Here, 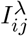 is 1 if the loop *Lij* is a *λ*-loop and 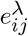 is the free energy of the loop *L*_*ij*_ when the loop *L*_*ij*_ is a *λ*-loop. Energy 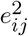 depends on the pairing positions *m* and *n* accessible from the positions *i* and *j*: 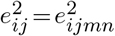 where 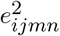 is the energy 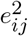 parameterized with the positions *m* and *n*. Turner’s model restricts the number of unpaired positions of the 2-loop *L*_*ij*_, (*m*−*i*)+(*j* −*n*)+2: (*m* − *i*)+(*j* − *n*)+2≤30 to reduce the time complexities of prediction algorithms. Energy 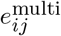 is decomposed into the terms of the closing and accessible pairings: 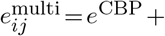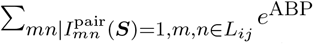 where *e*^CBP^ and *e*^ABP^ are free energy per closing and accessible pairing, respectively. Energy 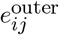 does not influence the entire free energy 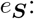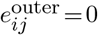:

As the components 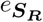 and 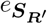, many conventional methods scoring structural alignments such as PMcomp, LocARNA, RAF, SPARSE, and LocARNA-P employ the *posterior model*. It scores the components 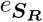 and 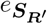 with posterior pairing probability matrices on structure (estimated by *inside-outside algorithm*s such as McCaskill’s algorithm (42) and its variant algorithms (43, 44, 45)) to simplify computations although the suitability of these matrices has not been discussed. (21, 26, 27, 29, 37) Hence, Turner’s model is adopted in this study to prevent a reduction in prediction accuracy due to a matrix 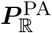.

The component 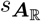 is computed by the learned parameters, including affine gap scores used in CONTRAlign, which predicts pairwise sequence alignments (46). The components *s*^PA^ and *s*^LA^ can be computed to sum RIBOSUM scores (1) across all pa aligned and loop-aligned positions, respectively: 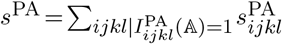 and 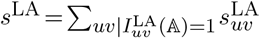 where 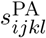 and 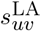 are the RIBOSUM pair and loop alignment scores of the pairs (*i,j*) and (*k,l*) and the positions *u* and *v*, respectively and 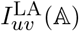 is 1 if the positions *u* and *v* are loop-aligned in the alignment 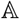 and 0 otherwise.

### Inside-outside algorithm that computes pair alignment probability matrix 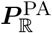

In this section, an efficient (however nevertheless impractical) method that computes a matrix 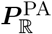 in the framework of inside-outside algorithm is proposed. A probability 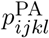 can be written with “inside” partition functions 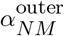 and 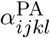 and an “outside” partition function 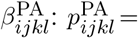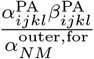 where *N* and *M* are the lengths of the sequences ***R*** and ***R***′, respectively. Inside and outside partition functions can be computed with those of shorter and longer substrings, respectively, and are stored in dynamic programming memory for the remaining computation. A matrix 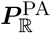 can be computed by Algorithm 1 with the *O*(*N*^4^*M*^4^) time and the *O*(*N*^3^*M*^3^) memory. (Figure 3a)

**Figure 3.**
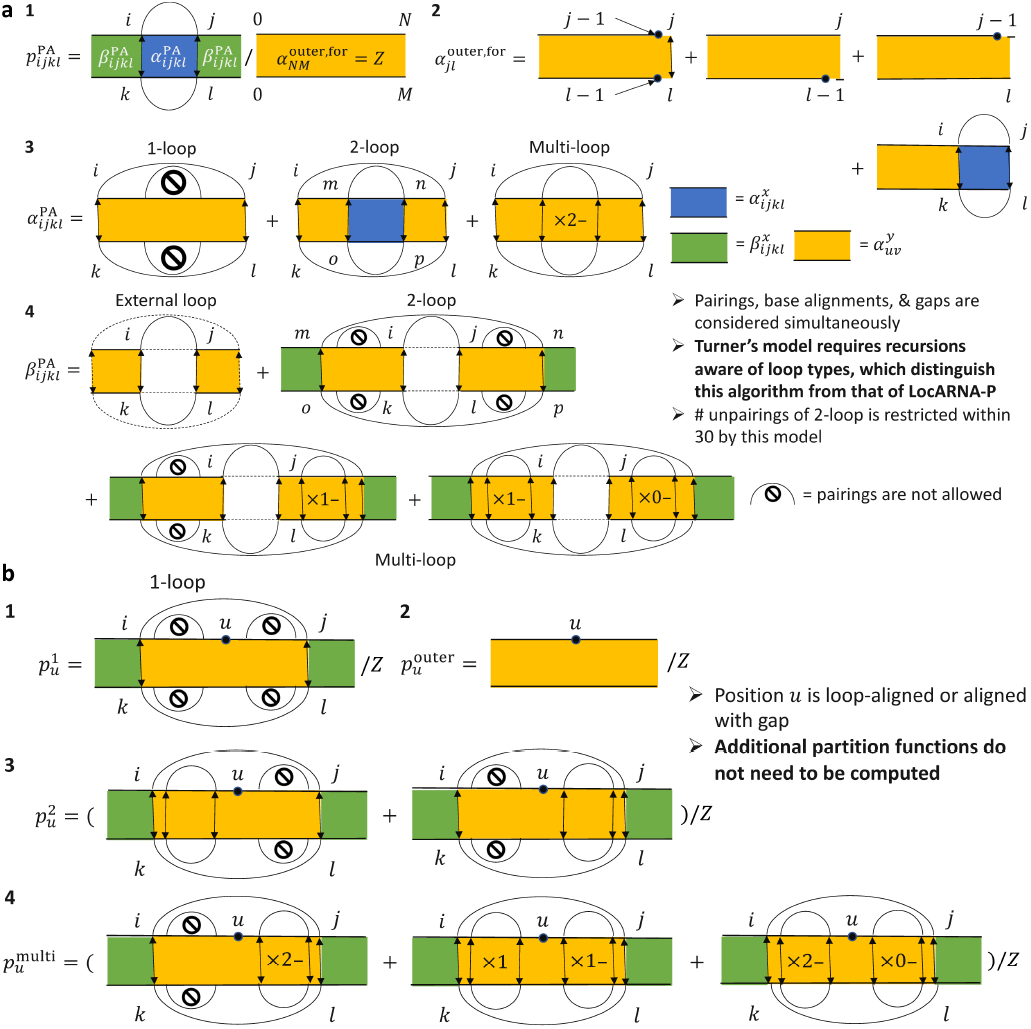
(a) An overview of recursions to compute (1) probabilities 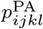 and partition functions (2) 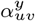, (3) 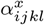, and (4) 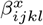, (b) An overview of recursions to compute accessibilities (1) 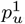, (2) 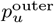, (3) 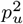, and (4) 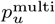.

**Algorithm 1.**
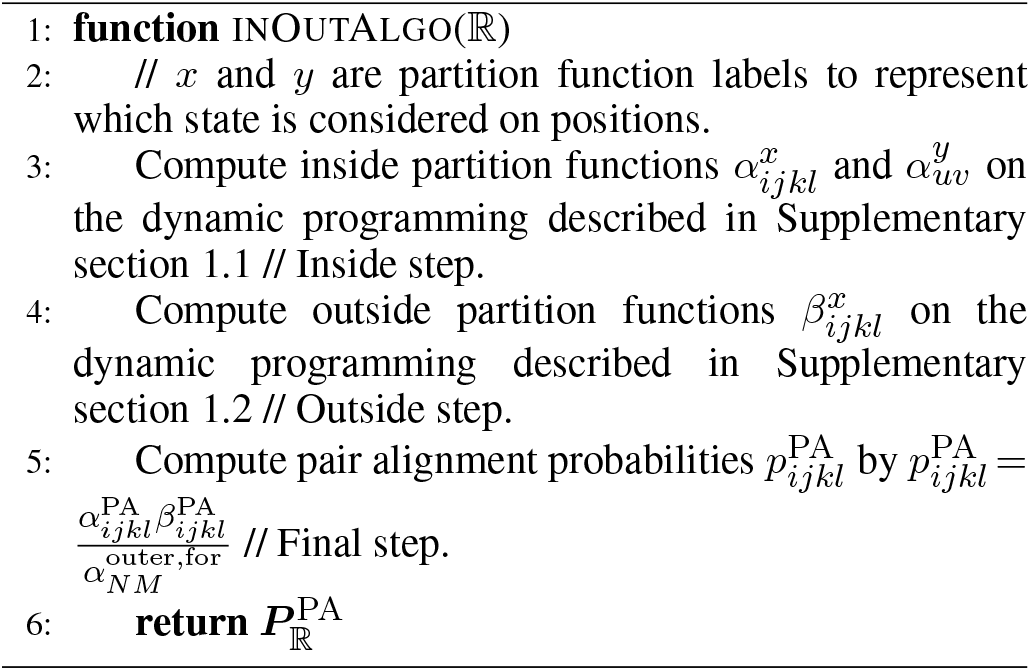
An inside-outside algorithm that computes a pair alignment probability matrix 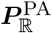

**Proof.**

Partition functions 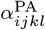 and 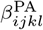 and probabilities 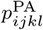 demand the *O*(*N*^2^*M*^2^) memory. Partition functions 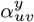 are stored for all the combinations of pair-aligned pairs (*i,j*) and (*k,l*) that close the positions *u* and *v*, respectively and thus demand the *O*(*N*^3^*M*^3^) memory.

Partition functions 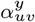 include the case where the positions *u* and *v* are pair-aligned and therefore demand the *O*(*N*^4^*M*^4^) time. Partition functions 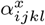 are computed from the partition functions 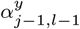 in only the *O*(1) time and thus demand the *O*(*N*^2^*M*^2^) time as a whole. Partition functions 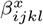 consider the pair-aligned pairs of positions that close the position pairs (*i,j*) and (*k,l*) and therefore demand the *O*(*N*^4^*M*^4^) time.

Probabilities 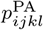 are computed from the partition functions 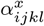 and 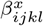 in only the *O*(1) time and therefore demand the *O*(*N*^2^*M*^2^) time as a whole. Finally, a matrix 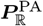 demands the *O*(*N*^4^*M*^4^) time and the *O*(*N*^3^*M*^3^) memory.

Algorithm 1 is the “simultaneous” solution of Durbin’s (*forward-backward*) algorithm, which estimates posterior alignment probability matrices on sequence alignment (47), and McCaskill’s algorithm, as expected. Algorithm 1 is also an inside-outside algorithm version of Sankoff’s algorithm, which predicts pairwise structural alignments whose score is maximum (18), as expected. However, desirable time and memory complexities of Algorithm 1 are quadratic to deal with long ncRNAs.

### Sparsifying pair alignment probability matrix 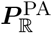

In this section, a solution to make Algorithm 1 lightweight, sparsifying all possible structural alignments 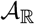, is introduced. From all the possible alignments 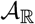, *sparsification* can pick out those favorable (e.g. with adequately high scores). It can allow Algorithm 1 to compute only the partition functions 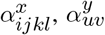, and 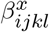 that satisfy sparsification conditions. (Figure 4) Let 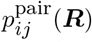 be the pairing probability of the positions *i* and *j* given the sequence ***R***. In this study, the following sparsification conditions are introduced:

- |*u*−*v*|≤*δ*^gap^ and |(*M*−*u*)−(*N*−*v*)≤*δ*^gap^ for any positions *u* and *v*
- |(*j*−*i*)−(*l*−*k*)|≤*δ*^gap^ for any pair-aligned pairs (*i,j*) and (*k,l*)
- 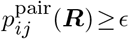 for any pairing positions *i* and *j* and any sequence ***R***

where *δ*^gap^ and *ϵ* are sparsification parameters. The first two *banding condition*s let Algorithm 1 not consider the alignments 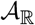 with too many gaps. (21, 22, 23, 24, 48)

**Figure 4.**
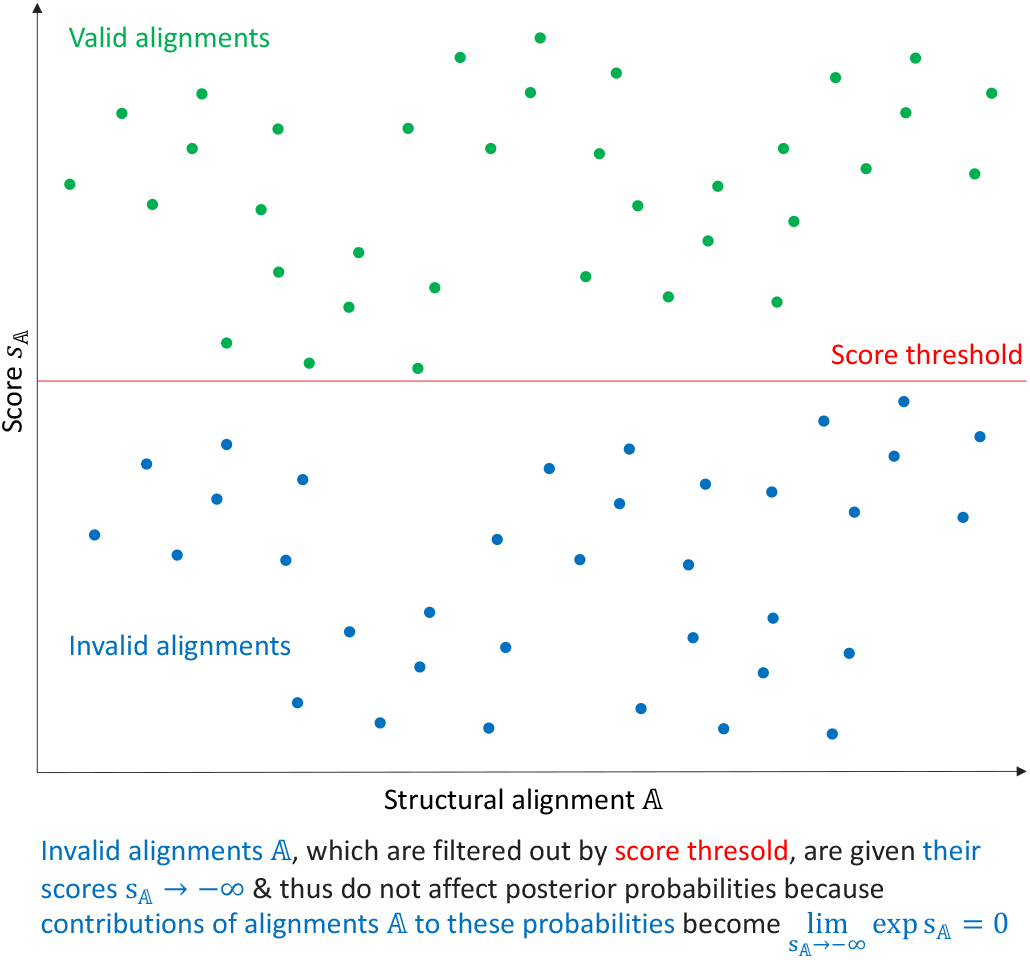
Sparsification lets Algorithm 1 ignore alignments 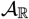 whose contributions to the partition function *Z* are low. Choosing sparsification conditions whose dynamic programming reduces effectively their computational complexities is of extreme importance.

The last *pairing condition* makes Algorithm 1 not consider the alignments 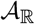 with pairings difficult to predict (e.g. distant). (24, 27, 28, 29, 37, 49, 50) If Turner’s model is replaced with the posterior model and the banding conditions are removed, Algorithm 1 becomes identical to LocARNA-P. (37) Algorithm 1 with the above conditions is ideal with the *O*(*L*^2^) time and the *O*(*L*^2^) memory where *L* = max(*N,M*) if the parameters *δ*^gap^ and *ϵ* take sufficiently small and large values, respectively.

**Proof.**

The numbers of all the possible pairings of positions *u* and *v* become *O*(*Nδ*^bp^) and *O*(*Mδ*^bp^) from *O*(*N*^2^) and *O*(*M*^2^), respectively where 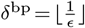 and [*r*] returns the greatest integer less than or equal to the real number *r*. The number of all the possible combinations of positions *u* and *v* becomes *O*(*Lδ*^gap^) from *O*(*NM*). The number of all the possible pair-aligned pairs (*i,j*) and (*k,l*) becomes *O*(#^PA^) from *O*(*N*^2^*M*^2^) where #^PA^ = *Lδ*^bp^*δ*^gap^*δ*^max^ and *δ*^max^ = max(*δ*^bp^*,δ*^gap^).

Partition functions 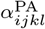 and 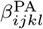 and probabilities 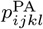 demand the *O*(#^PA^) memory. Partition functions 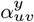 are stored for all the combinations of pair-aligned pairs (*i,j*) and (*k,l*) that close the positions *u* and *v*, respectively and thus demand the *O*(#^PA^*Lδ*^gap^) memory.

Partition functions 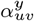 include the case where the positions *u* and *v* are pair-aligned and therefore demand the *O*((#^PA^)^2^) time. Partition functions 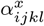 are computed from the partition functions 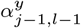 in only the *O*(1) time and thus demand the *O*(#^PA^) time as a whole. Partition functions 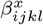 consider the pair-aligned pairs of positions that close the position pairs (*i,j*) and (*k,l*) and therefore demand the *O*((#^PA^)^2^) time.

Probabilities 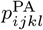 are computed from the partition functions 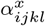 and 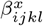 in only the *O*(1) time and therefore demand the *O*(#^PA^) time as a whole. Finally, a matrix 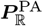 demands the *O*((#^PA^)^2^) time and the *O*(#^PA^*Lδ*^gap^) memory. If the parameters *δ*^gap^ and *δ*^bp^ are sufficiently small, a matrix 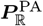 demands the *O*(*L*^2^) time and the *O*(*L*^2^) memory.

### Probabilistic consistency transformation

Probabilistic consistency transformation is the technique that converts a probability between a target homolog and each of its support homolog into a metric that summarizes the phylogeny among all the homologs. (24, 30, 31, 51) This transformation is required, because the computational complexities involved in computing posterior probabilities among all the homologs are NP-complete, as with multiple rigorous alignment. In this study, methods that transform probabilities 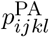 are proposed. To average probabilities 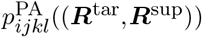 between the target homolog ***R***^tar^ and each support homolog ***R***^sup^ ∈ *R*^sup^, the average pairing 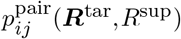 is gained:

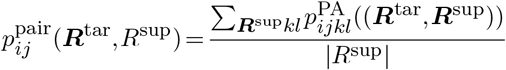

where *R*^sup^ is a set of support homologs of the homolog ***R***^tar^. To sum probabilities 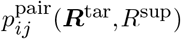, an average unpairing probability 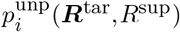 is obtained:

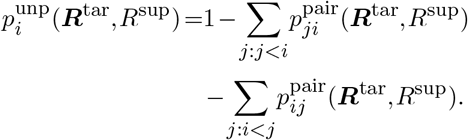

Obviously, the difference between proposed probabilities 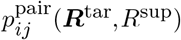 and existing probabilities 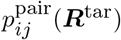 is whether support homologs *R*^sup^ are considered or not.

### Hom-folding that maximizes average expected accuracy

The accuracy of a predicted structure ***S*** against a reference alignment 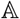 is measured based on terms of positive and negative predictions:

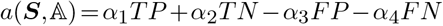

where *TP*, *TN*, *FP*, and *FN* are the numbers of true positive, true negative, false positive, and false negative predictions, respectively, and *α*_*h*_ are their scale parameters. In this study, the counts *TP*, *TN*, *FP*, and *FN* are configured as

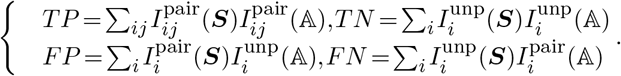

Here, 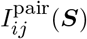 is 1 if the positions *i* and *j* are pairing in the structure ***S*** and 0 otherwise, 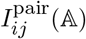 is 1 if the positions *i* and *j* are pairing in the alignment 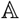 and 0 otherwise, 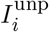 is 1 if the position *i* is unpaired in the structure ***S*** and 0 otherwise, 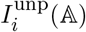 is 1 if the position *i* is unpaired in the alignment 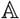 and 0 otherwise, 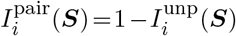, and 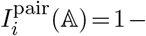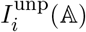

Because the accuracy 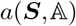 and the *γ*-dependent accuracy 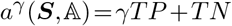 are equivalent, the expected accuracy to be maximized is gained:

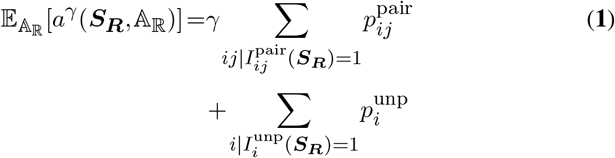

where 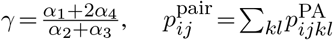, and 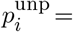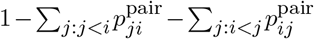

**Proof.**

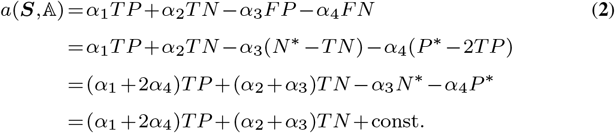

where 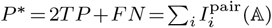 and 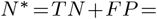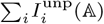. To divide the both sides of Equation **2** by the scaler *α*_2_+*α*_3_, the equivalence 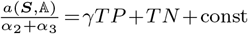. is obtained.

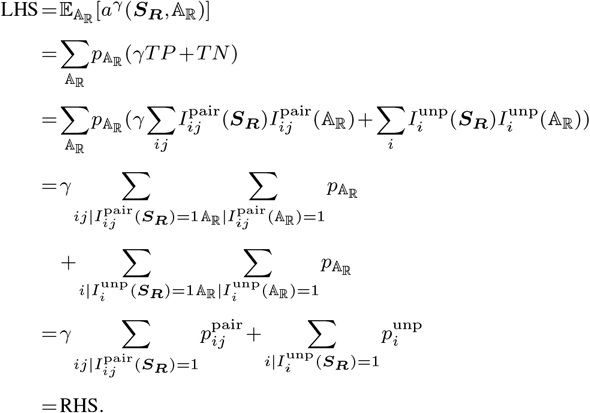

Deriving the expected accuracy that makes posterior probabilities explicit like Equation **1**, which enables dynamic programming to maximize this accuracy, is called *posterior decoding*. (52) The predicted structure that maximizes expected accuracy 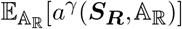 can be computed to conduct Nussinov-type dynamic programming (53) based on the following recursion:

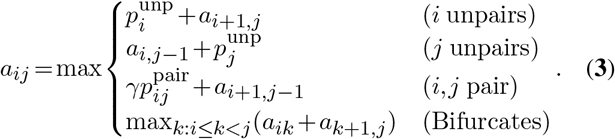

In Equation **3**, only the one support homolog ***R***′ of the target homolog ***R*** is considered to predict the single structure of the homolog ***R***. In order to consider more than one support homolog, it is sufficient that probabilities 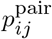 and 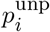 are replaced with the probabilities 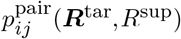 and 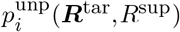 in Equation **3**, respectively. If the parameter *γ* is large, Equation **3** emphasizes positives and thus predicts more pairings. If the parameter *γ* is small, Equation **3** emphasizes negatives and thus predicts more unpairings.

### Ali-folding that maximizes mixed expected accuracy

Let 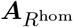 be the sequence alignment among the set of homologs *R*^hom^. Single structure prediction is extended to consensus structure prediction of sequence alignment to view positions *i* on a sequence ***R*** as columns *i* on an alignment 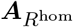 in Equation **3**. It is known that pairing probabilities of columns *i* and *j* given an alignment 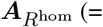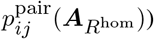, which can be computed by RNAalifold (32), improve the prediction accuracy of consensus structure. (32, 34) Thus, the mixture of probabilities 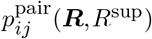 and 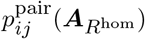 is used on Equation **3** to predict the consensus structure of an alignment 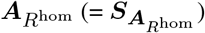:

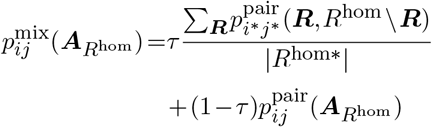

where 0≤*τ*≤1, ***R***∈*R*^hom^, *R*^hom*^ is the subset of the sequences *R*^hom^ that is not mapped to the gaps on the columns *i* and *j* in the alignment 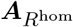, *i\** and *j\** are the positions on the sequence ***R*** mapped to the columns *i* and *j* in the alignment 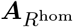 respectively. The parameter *τ* is a mixing coefficient. Likewise, the mixture of probabilities 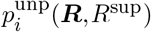 and 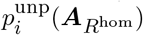 is used on Equation **3**:

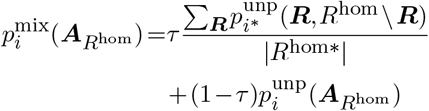

where 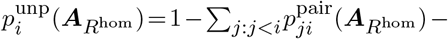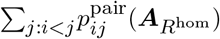. Finally, the following recursion, which predicts a structure 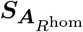, is obtained:

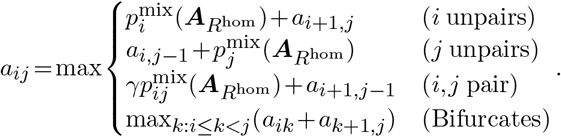

### Modifying Algorithm 1 to also compute average loop accessibilities on sparse pairwise structural alignment

The posterior probability that a position *u* is accessible from *λ*-loops is called the loop accessibility of the position *u*. Let loop accessibility matrices given a pair ℝ be 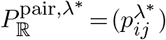 and 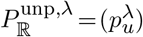 where *λ*\* ∈{2,multi,outer}, 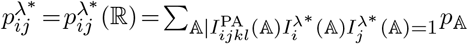 and 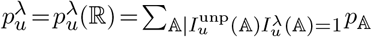. Here, 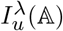 returns 1 if the position *u* is accessible from a *λ*-loop in the alignment 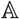. Accessibilities 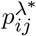 can be computed while computing probabilities 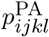 because 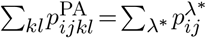. (Supplementary section 1.3) Accessibilities 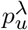 ask Algorithm 1 for additional computations. (Figure 3b) Matrices 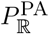, 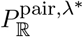 and 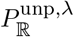 are computed by Algorithm 2 with the *O*(*N*^4^*M*^4^) time and the *O*(*N*^3^*M*^3^) memory. The sparsifications applied to Algorithm 1 are also applied to Algorithm 2. Therefore, the time and memory complexities of Algorithm 2 become *O*(*L*^2^) if the parameters *δ*^gap^ and *ϵ* take sufficiently small and large values, respectively as Algorithm 1.

Probabilistic consistency transformation for accessibilities 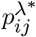 and 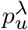 are also proposed. To average an accessibility 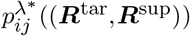 between the target homolog ***R***^tar^ and each support homolog ***R***^sup^, the average accessibility 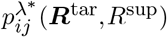 is obtained:

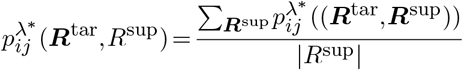

**Algorithm 2.**
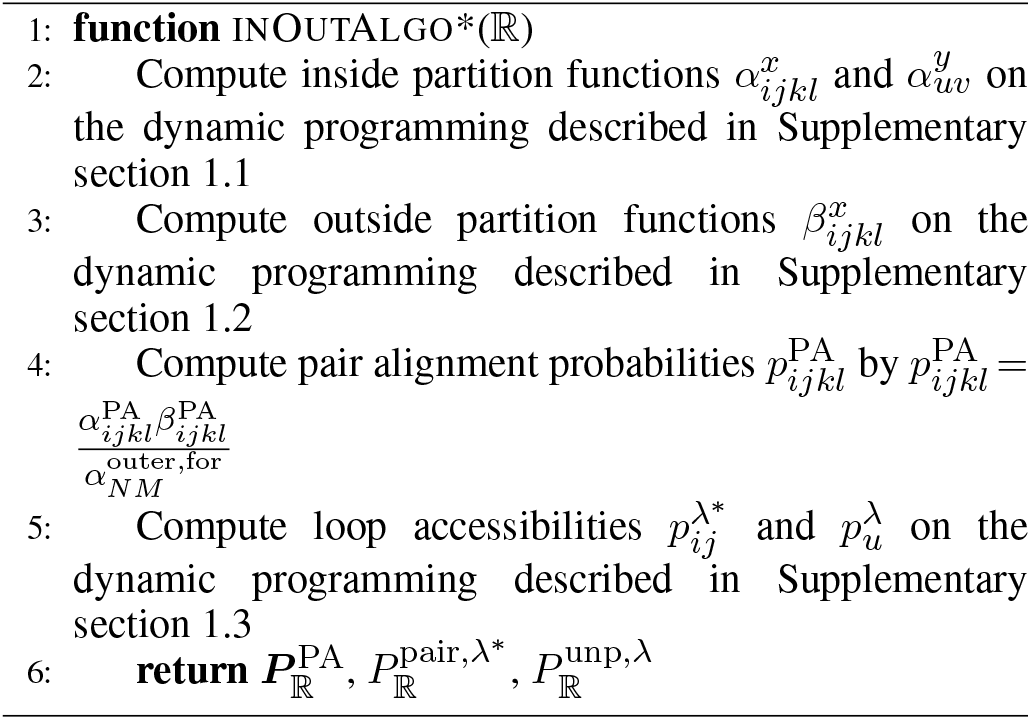
A variant algorithm of Algorithm 1 that computes a pair alignment probability matrix 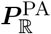 and loop accessibility matrices 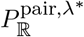 and 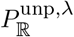

Likewise, the average accessibility 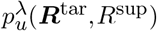 is gained:

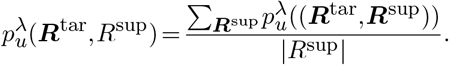

### Data collection for benchmark

From Rfam, which collects thousands RNA families (54), 1473 RNA families whose reference seed structural alignments had at most 200 columns and that contained at most ten sequences were collected as dataset “origin”. Reference single structures were obtained to map the reference seed consensus structure of each RNA family to each sequence on dataset “origin”. The obtained set which contains the sequences and their single structures of each RNA family is called test set “unaligned”. Reference consensus structures were obtained to leave only the reference seed consensus structure of each RNA family on dataset “origin”. The obtained set which contains the sequences and their consensus structure of each RNA family is called test set “aligned”.

### Competitors for benchmark

TurboFold v6.2, CentroidHomfold v0.0.16, CONTRAfold v2.02, CentroidFold v0.0.16, and RNAfold v2.4.14 (Table 1) were compared to ConsHomfold using their default parameters.

**Table 1.**
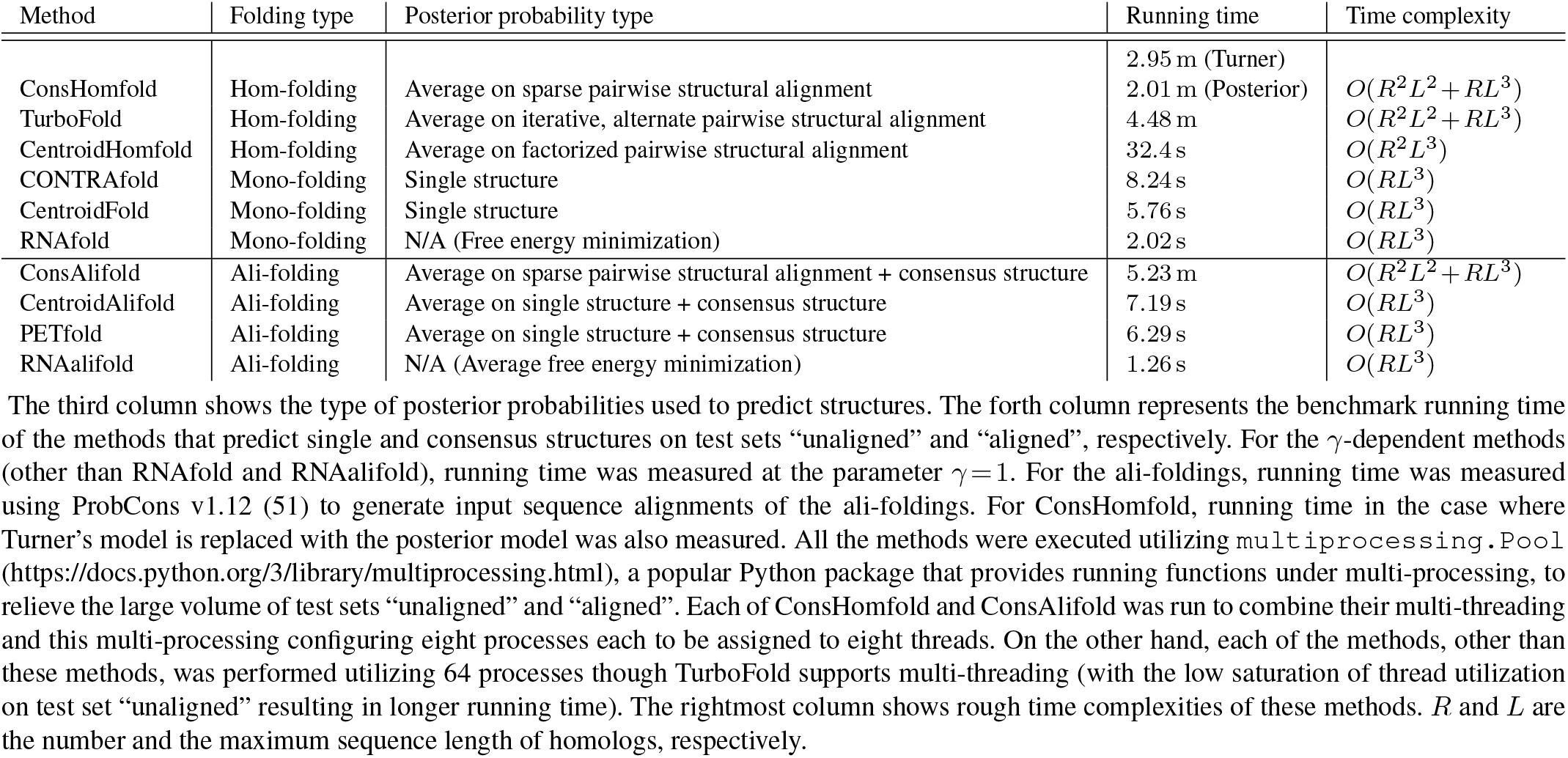
The profile and benchmark running time of methods that predict structures.

*RNAfold* RNAfold is the most standard mono-folding and predicts a structure ***S***_***R***_ to minimize its energy: 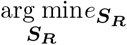. (4) This minimization is equivalent to the maximum likelihood prediction 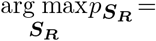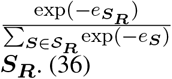 where ***S***_***R***_ is a set of all possible structures ***S***_***R·***_ (36)

*CONTRAfold* CONTRAfold is a mono-folding that predicts a structure ***S***_***R***_ maximizing its expected accuracy

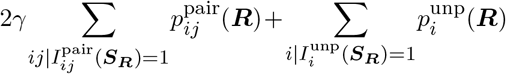

where 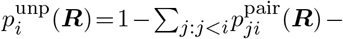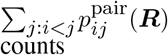. (8, 9) This accuracy is based on the counts

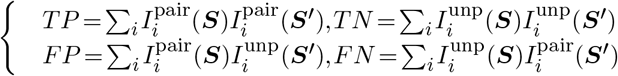

where ***S***′∈***S***_***R***_. The count *TP* does not mind the pairing partner *jpairing with at most one position* of a position *i* if only the position *i* is pairing.

*CentroidFold* CentroidFold is a mono-folding that predicts a structure ***S***_***R***_ maximizing its expected accuracy 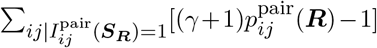. (9) This accuracy is based on the counts

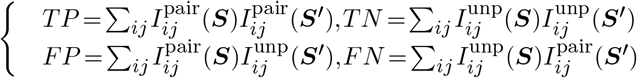

where 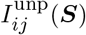 is 1 if the positions *i* and *j* are unpairing in the structure ***S*** and 0 otherwise. The counts *TP*, *TN*, *FP*, and *FN* are biased to negatives since a position *i* can be pairing with at most one position *j* and thus most pairs (*i,j*) are unpairing. This bias becomes remarkable when sequences are long.

*TurboFold* TurboFold is a hom-folding that iteratively, alternately estimates posterior probabilities on single structure and those on pairwise sequence alignment of homologs. (31) During this estimation (called the *turbo decoding* (55, 56)), the probabilities estimated currently (e.g. on pairwise sequence alignment) are incorporated into those estimated immediately afterwards (e.g. on single structure). After *η* iterations of the alternate estimation, TurboFold predicts the single structures of the homologs to maximize the expected accuracy based on the posterior probabilities estimated finally by the turbo decoding. From the viewpoint of structural alignment, TurboFold is said to decompose a multiple structural alignment of homologs into its single structures and pairwise sequence alignments on probability. TurboFold predicts a structure 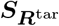 maximizing its expected accuracy

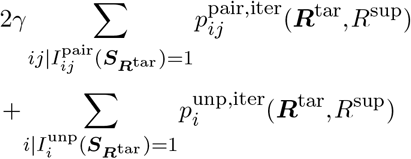

where 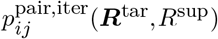 is the average pairing probability of the positions *i* and *j* given the sequence ***R***^tar^ and the sequences *R*^sup^ estimated by the turbo decoding and 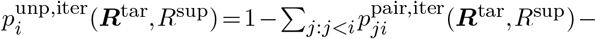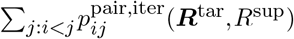. In the benchmark of this study, TurboFold was retried with the parameter *η* = 1 when this method failed with its default parameters (including the parameter *η* = 3).

*CentroidHomfold* CentroidHomfold is a hom-folding that extends CentroidFold to incorporate homologs and *factorize*s a probability 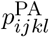 into independent posterior probabilities:

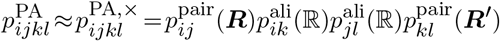

where 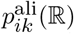 is the alignment probability of the positions *i* and *k* given the pair ℝ on pairwise sequence alignment. This factorization lets CentroidHomfold avoid the computation of probabilities 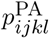. (30) CentroidHomfold connects probabilities 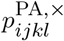 into a single metric via probabilistic consistency transformation as ConsHomfold and then predict the single structures of the homologs using the metrics obtained by this transformation. CentroidHomfold predicts a structure 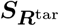 maximizing its expected accuracy

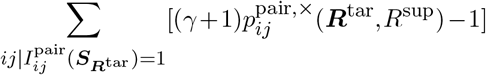

where 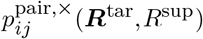 is the average pairing probability of the positions *i* and *j* given the sequence ***R***^tar^ and the sequences ***R***^sup^ obtained by this transformation.

CentroidAlifold v0.0.16, RNAalifold v2.4.14, and PETfold v2.1 (Table 1) were compared to ConsAlifold using their default parameters.

*RNAalifold* RNAalifold is the most standard ali-folding and predicts a structure 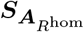 minimizing its average energy 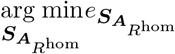 where 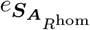 is the average energy of the structure 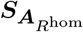. (32) This minimization is equivalent to the maximum likelihood prediction 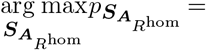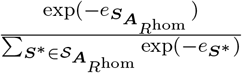 where 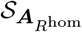 is a set of all possible structures 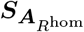. (36)

*CentroidAlifold* CentroidAlifold is an ali-folding that predicts a structure 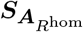 maximizing its expected accuracy 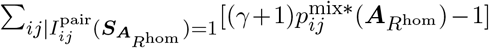 where 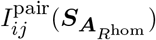 1 if the columns *i* and *j* are pairing in the structure 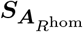 and 0 otherwise and 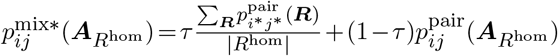.(34)

*PETfold* PETfold is an ali-folding that predicts a structure 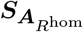 to maximize its expected accuracy

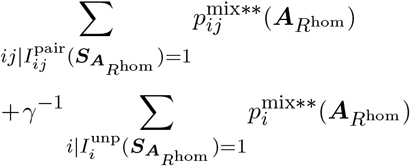

where 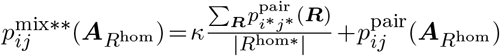, 0<*k*, 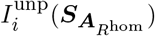 returns 1 if the column *i* is unpairing in the structure 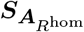 and 0 otherwise, and 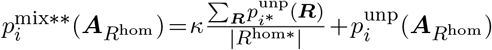. (33) CentroidAlifold and PETFold become equivalent to McCaskill-MEA (57) when *τ* = 1 and *κ*→∞ except their count configuration, respectively.

### CapR

CapR v1.1.1 was compared to ConsHomfold and ConsAlifold. CapR computes loop accessibilities

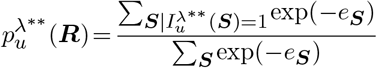

where *λ*\*\* ∈{1,stem,bulge,interior,multi,outer} and 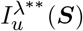 returns 1 if the position *u* is accessible from a *λ*\*\*-loop in the structure ***S***. (39) Here, the position *u* is said to be in a stem loop if the position *u* closes a stacking or is accessible from it. Also, CapR changes the definition of a (*λ*\*\* ≠ stem)-loop to exclude pairing positions in the (*λ*\*\* ≠ stem)-loop from this definition. To enable genomewide analysis, this method considers all possible local structures ***S*** |*j*−*i*|≤*W* on any pairing pairs *i*,*j*) where *W* is the *maximum span* of pairings, which regulates the structures ***S*** (43). In this study, the span *W* = 200 was used though the length of the sequence applied CapR to was less than 200, i.e. this method took all possible structures ***S*** into account. CapR does not incorporate support homologs of a target homolog.

### Metrics for prediction accuracy

Positive predictive value (= PPV), sensitivity, false positive rate (= FPR), the F1 score, and the Matthews correlation coefficient (= MCC) are calculated from the numbers of true and false positives and negatives:

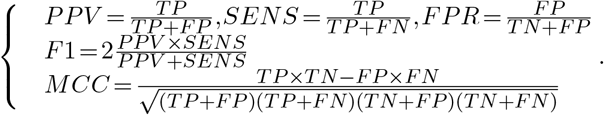

For single structures, the counts *TP*, *TN*, *FP*, and *FN* are configured as

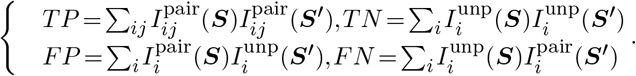

For consensus structures, the counts *TP*, *TN*, *FP*, and *FN* are configured in two ways as

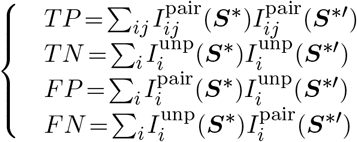

(called *columnwise count*s) where 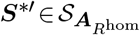 and

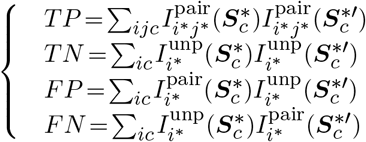

(called *mapwise count*s) where 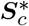 and 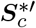 are the single structures obtained to map the structures ***S***\* and ***S***\*′ to the *c*-th sequence, respectively.

### Implementations and benchmark environments

ConsHomfold and ConsAlifold implemented in Rust employ multi-threading to give their users more efficient computing. Probabilities and partition functions are computed under the log scale using the logsumexp trick *log*∑_*a*_*expx*_*a*_=*log*∑_*a*_*exp*(*x*_*a*_−*max*_*a*_))+*max*_*a*_, which mitigates the undesirable effect of extremely large and small values (e.g. the overflow and underflow of floating point values), in these methods where *x*_*a*_ is a real number. Sparse data structures were implemented by `FxHashMap` (the fastest, memoryefficient hash table in Rust to our best knowledge) provided by https://github.com/Amanieu/hashbrown. The implementation choice of these structures is critical because the efficiency of these structures dominates the entire running time and memory usage of the methods. In this study, the methods used the parameters

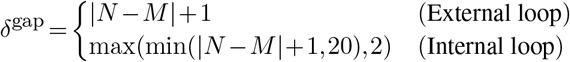

*ϵ* = 0.005, and *τ* = 0.5 (used as the default value of the parameter *τ* by CentroidAlifold (34)). For the comparison with Turner’s model, the posterior model was also implemented in ConsHomfold to use the scoring 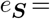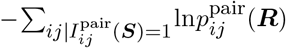. For the benchmark of running time, programs were run on a computer composed of an “Intel Xeon CPU” CPU with 64 threads and a clock rate of 2.30 GHz and 240 GB of RAM. Otherwise, programs were run on a computer composed of an “AMD EPYC 7501” CPU with 64 threads and a clock rate of 2 GHz and 128 GB of RAM.

## RESULTS

### Benchmark of ConsHomfold and ConsAlifold with their competitors

ConsHomfold and ConsAlifold perform the best trade-off of the metrics *PPV*, *SENS*, and *FPR* (Figure 5a) among the state-of-the-art methods that predict structures while requiring comparable running time (Table 1). Also, ConsAlifold demonstrates better transitions of the metrics *F* 1 and *MCC* than CentroidAlifold and PETfold. (Figure 5b3–4) However, PETfold does not drop the metrics *F* 1 and *MCC* within the range −7≤log_2_*γ*≤−4 compared to ConsAlifold and CentroidAlifold. (Figure 5b3–4) The above accuracy performances on consensus structure are also confirmed when columnwise counts were used instead of those mapwise. (Figure 6) RNAalifold shows competitive accuracy on all the metrics *PPV*, *SENS*, *FPR*, *F* 1, and *MCC*, except the metric *MCC* when ProbCons was used. (Figure 5ab, Figure 6) The columnwise counts suffer from the quality of input sequence alignments compared to those mapwise. (Figure 5ab, Figure 6)

**Figure 5.**
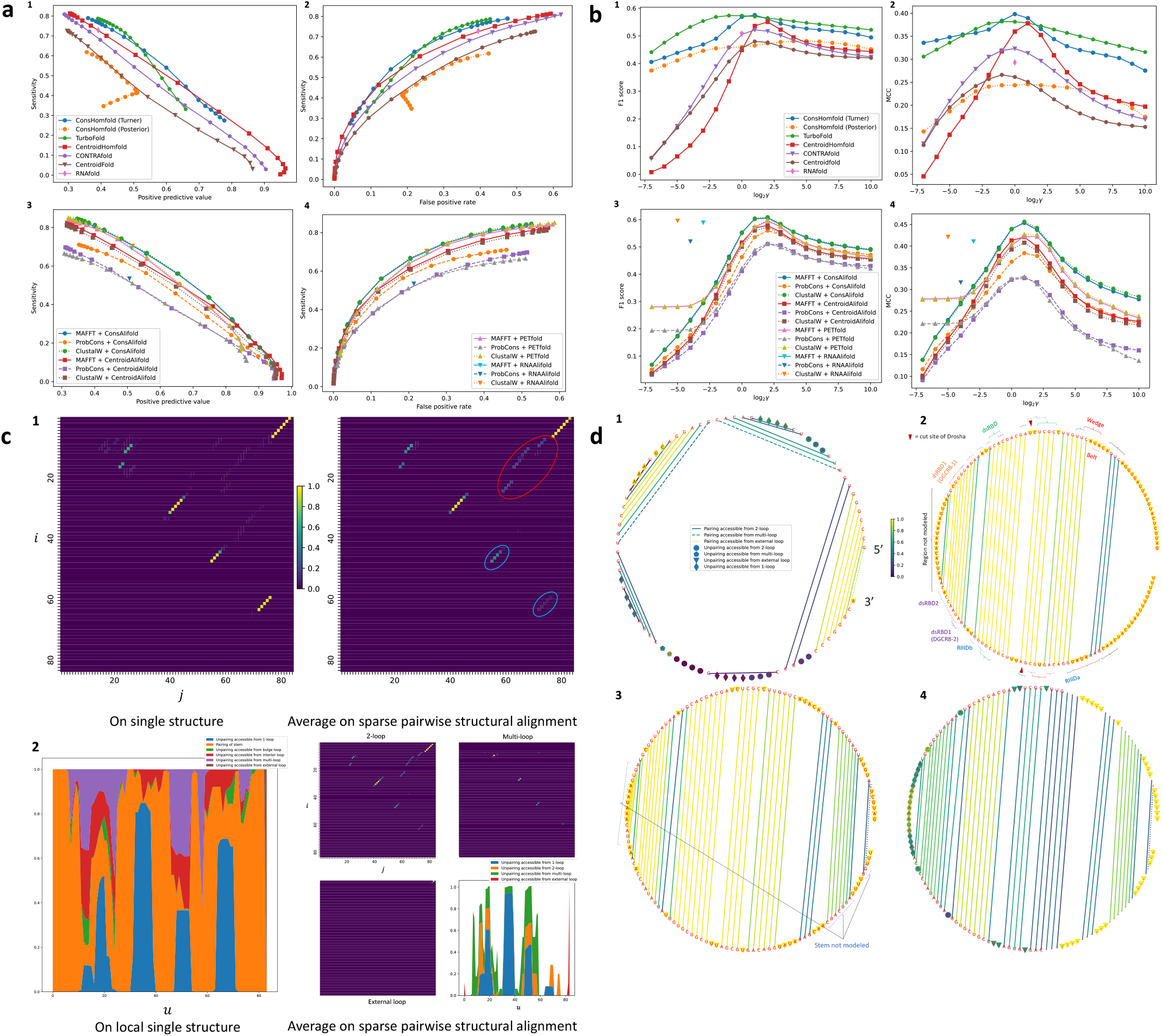
**(a) The trade-off curves composed of (1, 3) pairs** (*PPV,SENS*) **and (2, 4) pairs** (*FPR,SENS*) **at each parameter** *γ* = 2^*g*^:*g*∈{−7,…,10, **respectively.** These curves are unavailable for RNAfold and RNAalifold because they do not depend on the parameter *γ*. The methods that predict (1, 2) single and (3, 4) consensus structures were measured using test sets “unaligned” and “aligned”, respectively. Mapwise counts were used for consensus structures. Turner’s and the posterior models were compared on ConsHomfold. The MAFFT v7.470 (61), ProbCons, and ClustalW v2.1 (70) were used with their default parameters to generate input sequence alignments of the ali-foldings. **(b) The transitions of the metrics (1, 3)***F* 1 **and (2, 4)***MCC* **across parameters** *g* = log_2_ *γ***, respectively**. The used test sets, the comparison between Turner’s and posterior models, counts for consensus structures, and input alignment settings are the same as Figure 5a. For RNAfold, the metrics *F* 1 and *MCC* are plotted at the parameter *g* = 0. For RNAalifold, the metrics *F* 1 and *MCC* computed with MAFFT, ProbCons, and ClustalW are plotted at the parameters *g* = −3, *g* = −4, and *g* = −5, respectively. **(c) The various probability distributions of a tRNA.**(1) The comparison between probabilities (left) 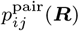 and (right) 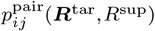. Probabilities 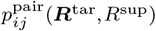 are supported by other five tRNAs. The red and blue circles display conserved and nonconserved pairings, respectively. (2) The comparison between (left) accessibilities 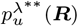 and (right) accessibilities 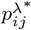 and 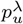. **(d) The structures of a tRNA and pri-miR-16-2 color-coded by accessibilities** 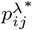 **and** 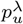. (1) The single structure of a tRNA supported by other five tRNAs predicted by ConsHomfold at the parameter *γ* = 2^10^ = 1024. The single structures of pri-miR-16-2 (2) modeled in the study that determined the Cryo-EM structure of human Drosha and DGCR8 binding this RNA (60) and (3) predicted by ConsHomfold at the parameter *γ* = 2^3^ = 8 using the ten homologs of this RNA. Each pairing and unpairing are color-coded by maximum accessibilities 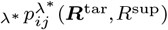 and 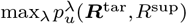. The sites of this model structure bound by components of Drosha and DGCR8 retrieved from the study are also displayed. Belt, Wedge, dsRBD, RIIIDa, and RIIIDb are components of Drosha. dsRBD1 and dsRBD2 are components of DGCR8. DGCR8-1 and DGCR8-2 are different copies of DGCR8. (4) The consensus structure predicted by ConsAlifold at the parameter *γ* = 2^3^ = 8. This structure is of the sequence alignment among pri-miR-16-2 and the homologs of this RNA predicted by MAFFT with its default parameters. Each column of this alignment is represented by the most frequent character including a gap in this column. Each pairing and unpairing are color-coded by maximum accessibilities 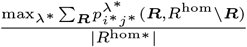 and 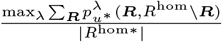 where *u*\* is the position on the sequence ***R*** mapped to the column *u* in this alignment.

**Figure 6.**
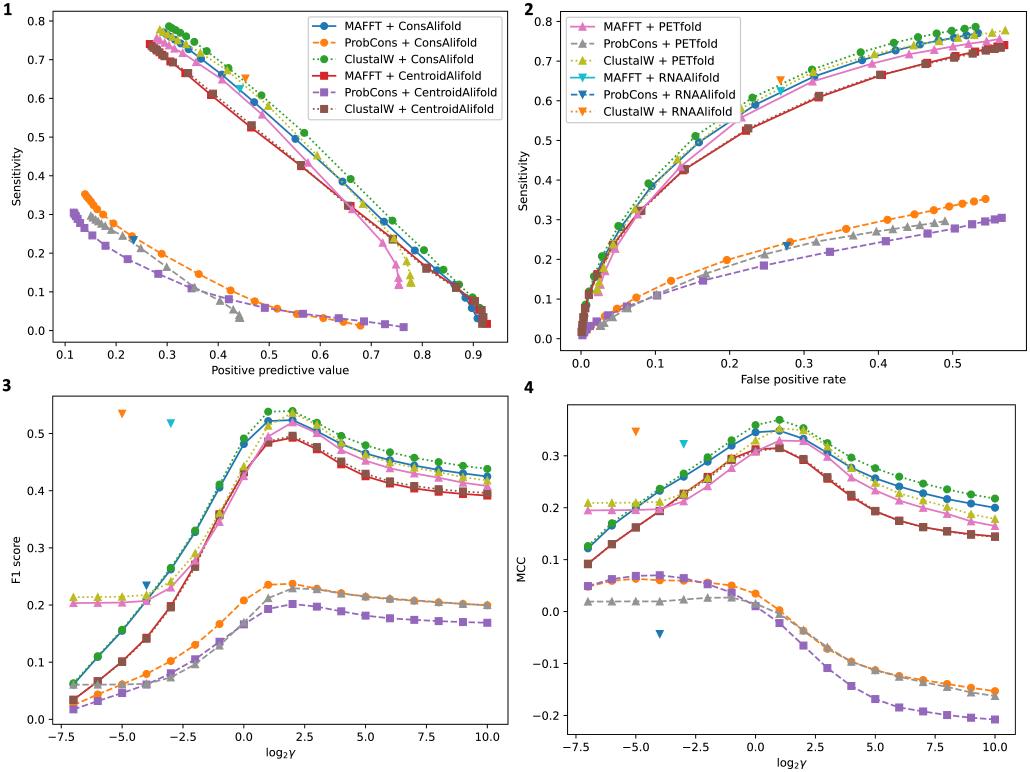
The trade-off curves composed of (1) pairs (*PPV,SENS*) and (2) pairs (*FPR,SENS*) at each parameter *γ* = 2^*g*^ on columnwise counts, respectively. (b) The transitions of the metrics (3) *F* 1 and (4) *MCC* across parameters *g* = log_2_ *γ* on columnwise counts, respectively. The ali-foldings were measured using test set “aligned”. The input alignment settings are the same as Figure 5a.

TurboFold displays a superior transition of the metric *F* 1 to the other methods that predict single structures whereas competing with ConsHomfold on the metric *MCC*. (Figure 5b1–2) As expected, ConsHomfold with the posterior model exerts significantly less predictive power than that with Turner’s model across all the metrics *PPV*, *SENS*, *FPR*, *F* 1, and *MCC* (Figure 5ab), though the former records the faster running time than the latter (Table 1).

### Comparison between conventional and proposed posterior probabilities of example ncRNA: tRNA

The conventional probabilities 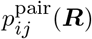 and accessibilities 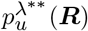 of a tRNA are contrasted to its proposed probabilities 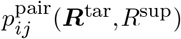 and accessibilities 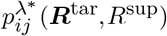 and 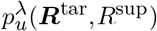 in terms of homology consideration. (Figure 5c) Conserved and nonconserved pairings are clarified through the gaps between the probabilities 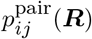 and 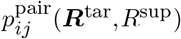 of this RNA. (Figure 5c1)

### Proposed loop accessibilities of example ncRNAs: tRNA and microRNA

The single (cloverleaf) structure of a tRNA predicted by ConsHomfold is decorated with the proposed accessibilities 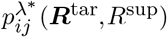 and 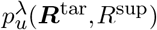. (Figure 5d1) The reliability of each pairing and unpairing in this structure can be assessed through the accessibilities 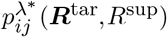 and 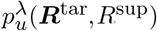.

The single structure of pri-miR-16-2, which is one of primary microRNAs (= pri-miRNAs), modeled to investigate the mechanism of human Drosha and DGCR8 (58) that cleaves metazoan pri-miRNAs in order to generate their mature miRNAs through Dicer (59) using Cryo-EM (60) is compared to that predicted by ConsHomfold. (Figure 5d2–3) The outlines of these structures agree though stems not found in the model structure were mispredicted by ConsHomfold. (Figure 5d2–3) The reliability of these stems is as high as the other parts of the predicted structure. (Figure 5d3) Accessibilities 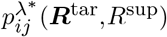 and 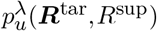 can help biologists validate model single structures as in the presented model structure. The entire reliability of the consensus structure of pri-miR-16-2 and its homologs predicted by MAFFT and ConsAlifold is lower than that of the structure predicted by ConsHomfold though the contours of these structures coincide. (Figure 5d3–4)

## DISCUSSION

### Probabilistic, consistent consideration is possibly simple answer for further improvement of prediction accuracy

Considering consistently pairwise structural alignments on structure prediction is the prospect to improve further this prediction, though the effectiveness of this consideration has not been focused on. ConsHomfold and ConsAlifold demonstrate that this consideration improves the prediction accuracy on this prediction. This improvement is possibly successful in resolving other prediction problems, such as sequence alignment predictions and certainly structural alignment predictions. It is likely that CentroidAlign (49) and MAFFT (61), which can predict sequence alignments considering structural alignments via their decomposition, will become universally accepted for the adoption of the consistent consideration, instead of this decomposed consideration. DAFS, which can predict structural alignments with this decomposed consideration (28), will be also enhanced by this adoption.

### Turner’s model is also effective in structural alignments

A majority of conventional methods that predict structural alignments use the posterior model to score the single structures in structural alignments being computed, because aligning more than two sequences with Turner’s model (expected to display more predictive power than the posterior model) is computationally complicated. However, Turner’s model can be used avoiding this complication to predict structural alignments of the maximum expected accuracy principle as DAFS (28) where probabilistic consistency transformation, which decomposes (NP-complete) multiple structural alignments to be considered into pairwise structural alignments, is available. It is valuable that rebuilding popular methods that predict structural alignments, such as LocARNA and SPARSE, in this principle on Turner’s model, because ConsHomfold proves that this model is superior to the posterior model in terms of prediction accuracy.

### Extending ConsHomfold and ConsAlifold to more sophisticated prediction of structures

At present, ConsHomfold and ConsAlifold cannot predict pseudoknotted structures and those enhanced by single structure probing data. Augmenting these methods to also predict these structures is a straightforward future task that can be explored in the future. An algorithm that computes posterior pairing probabilities on pseudoknotted single structure (62, 63) cannot be simply extended to compute those on pseudoknotted pairwise structural alignment, because this algorithm demands the *O*(*N*^5^) running time and the *O*(*N*^4^) memory usage (41) whereas McCaskill’s algorithm demands the *O*(*N*^3^) running time and the *O*(*N*^2^) memory usage (42). IPknot offers a reasonable way to use McCaskill’s algorithm for the prediction of pseudoknotted structures. (41) More specifically, this method decomposes a pseudoknotted structure into its pseudoknot-free structures and then maximize the expected accuracy of these structures based on posterior pairing probabilities on pseudoknot-free single structure computed by McCaskill’s algorithm. Importing the methodology of IPknot and replacing McCaskill’s algorithm with Algorithm 1 and the proposed probabilistic consistency transformation in this methodology, ConsHomfold and ConsAlifold will predict pseudoknotted structures.

RNAfold and RNAalifold can predict structures limited by SHAPE reactivity data. (64) TurboFold can conduct its hom-folding utilizing this data. (65) Many methods that predict structures incorporating the data including the above methods add a new term called *SHAPE-origin pseudo-free energy* to the free energy of a structure. (64, 65, 66, 67, 68, 69) This pseudo-energy is obtained to convert the data into the pseudo-free energy per unpairing position and then sum this energy across all unpairing positions. ConsHomfold and ConsAlifold will predict structures constrained by the data to introduce SHAPE-origin pseudo-free energy.

### Proposed loop accessibilities are worth using in field of RNA-binding protein

CapR revealed that several RNA-binding proteins bind their target RNAs by recognizing the loop types of the binding regions in the RNAs (71, 72, 73) through CLIP-seq data (74). (39) The proposed loop accessibilities in this study can be used to analyze these proteins and the target RNAs including their structures more precisely. RNAcontext (75) and RCK (76) demonstrated that loop accessibilities improve the prediction accuracy of predicting the binding of the proteins to their candidate target RNAs compared to the conventional methods that do not use these accessibilities, such as MEMERIS (77) and MatrixREDUCE (78). The proposed accessibilities have the potential to ameliorate the prediction accuracy of RNAcontext and RCK.

## CONCLUSION

In this study, the below approaches have been proposed:

- a hom-folding (ConsHomfold) and an ali-folding (ConsAlifold) that consider sparse pairwise structural alignments on their probability distributions
- a quadratic homolog-aware algorithm to compute different kinds of average posterior probabilities on sparse pairwise structural alignment, some of which are used to conduct these foldings, and the others of which are helpful in analyzing RNAs and their structures.

ConsHomfold and ConsAlifold have represented better trade-off between PPV, sensitivity, and FPR on Rfam-based benchmarks than other state-of-the-art methods that predict structures including those that consider stochastically pairwise structural alignments to decompose them into their independent components. ConsAlifold has also displayed superior transitions of the F1 score and the MCC to the conventional methods of this method. From these results, It has been concluded that the consistent, probabilistic consideration of sparse pairwise structural alignments improves the prediction accuracy of structures.

ConsHomfold and ConsAlifold demand the reasonable running time supported by the compared time complexities of the benchmarked methods. It has been confirmed that Turner’s model, which is the most popular to score single structures, adopted in this study significantly more fits to score structural alignments than the posterior model which is used by many conventional methods that score structural alignments. Turner’s model possibly raises the prediction accuracy of methods that predict structural alignments using the posterior model.

Conventional loop accessibilities on single structure succeeded in the analysis and prediction of RNA-binding proteins and the *in silico* RNA aptamer selection on HT-SELEX (79, 80). There is a possibility that the loop accessibilities proposed in this study have a broader range of impactful applications, as with the conventional accessibilities.

## DATA AVAILABILITY

ConsHomfold, ConsAlifold, the data used in this study, and Python scripts to generate the figures and tables in this study are freely available at https://github.com/heartsh/conshomfold and https://github.com/heartsh/consalifold managed by M.T.

## SUPPLEMENTARY DATA

Supplementary Data is available from NAR Online.

## FUNDING

This work was not supported by any funding.

## Conflict of interest statement

None declared.

## ACKNOWLEDGEMENTS

We would like to thank Dr. Kiyoshi Asai, Dr. Martin Frith, and the members of their laboratories for discussing this study with us for years. Also, we offer our thanks to Dr. Risa Kawaguchi for sharing valuable information about the scaling and logsumexp methods on exact probability computations with us. We are appreciate being advised on work closely related to this study, LocARNA-P and PETfold, by kind anonymous reviewers. We would like to thank Editage (www.editage.com) for English language editing. Most computations were performed on the NIG supercomputer at the ROIS National Institute of Genetics.

## AUTHOR CONTRIBUTIONS

M.T. conceived the proposed method, implemented it in the presented programs, experimented with them, and wrote this paper.

## Notes

### Competing Interest Statement

The authors have declared no competing interest.

