## Supplementary materials for "Consistent Consideration of RNA Structural Alignments Improves Prediction Accuracy of RNA Secondary Structures"

### 1 Recursions of partition functions $\alpha^{x}_{ijkl}$, $\alpha^{y}_{uv}$, and $\beta^{x}_{ijkl}$ for Algorithm 1

#### 1.1 Recursions for efficient computations of inside partition functions $\alpha^{x}_{ijkl}$ and $\alpha^{y}_{uv}$

Let $\alpha^{\mathrm{PA}}_{ijkl}$ be the inside partition function between the pair-aligned pairs $(i,j)$ and $(k,l)$. A partition function $\alpha^{\mathrm{PA}}_{ijkl}$ consists of three additional components: $\alpha^{\mathrm{PA}}_{ijkl} = \alpha^{\mathrm{PA},1}_{ijkl} + \alpha^{\mathrm{PA},2}_{ijkl} + \alpha^{\mathrm{PA,multi}}_{ijkl}$ where $\alpha^{\mathrm{PA},1}_{ijkl}$, $\alpha^{\mathrm{PA},2}_{ijkl}$, and $\alpha^{\mathrm{PA,multi}}_{ijkl}$ are the partition function $\alpha^{\mathrm{PA}}_{ijkl}$ whose loops $L_{ij}$ and $L_{kl}$ are 1-loops, 2-loops, and multi-loops, respectively. The first term $\alpha^{\mathrm{PA},1}_{ijkl}$ recurses into its inner partition function: $\alpha^{\mathrm{PA},1}_{ijkl} = \exp(s^{\mathrm{ali}}_{ik} + s^{\mathrm{ali}}_{jl} + s^{\mathrm{PA}}_{ijkl} - e^{1}_{ij} - e^{1}_{kl})\alpha^{\mathrm{SA,for},(i,j),(k,l)}_{j-1,l-1}$ where $s^{\mathrm{ali}}_{ik}$ is the CONTRAlign emission score of the aligned positions $i$ and $k$ and $\alpha^{\mathrm{SA,for},(i,j),(k,l)}_{np} = \alpha^{\mathrm{SA,for}}_{np}$ is the partition function between the pairs $(i+1,n)$ and $(k+1,p)$ whose loops $L_{ij}$ and $L_{kl}$ are 1-loops. A partition function $\alpha^{\mathrm{SA,for}}_{np}$ is composed of three terms: $\alpha^{\mathrm{SA,for}}_{np} = \alpha^{\mathrm{SA,for,ali}}_{np} + \alpha^{\mathrm{SA,for,gap}}_{np} + \alpha^{\mathrm{SA,for,gap}\prime}_{np}$ where $\alpha^{\mathrm{SA,for,ali}}_{np}$, $\alpha^{\mathrm{SA,for,gap}}_{np}$, and $\alpha^{\mathrm{SA,for,gap}\prime}_{np}$ are the partition function $\alpha^{\mathrm{SA,for}}_{np}$ in the cases in which the positions $n$ and $p$ are aligned, the position $n$ is aligned with a gap immediately behind the position $p$, and the position $p$ is aligned with a gap immediately behind the position $n$, respectively. Partition functions $\alpha^{\mathrm{SA,for,ali}}_{np}$, $\alpha^{\mathrm{SA,for,gap}}_{np}$, and $\alpha^{\mathrm{SA,for,gap}\prime}_{np}$ recurse into their forward partition functions:

$$\begin{cases} \alpha^{\mathrm{SA,for,ali}}_{np} = (\alpha^{\mathrm{SA,for,ali}}_{n-1,p-1}\exp(s^{\mathrm{ali}\to\mathrm{ali}}) + \alpha^{\mathrm{SA,for,gap}}_{n-1,p-1}\exp(s^{\mathrm{gap}\to\mathrm{ali}}) + \alpha^{\mathrm{SA,for,gap}\prime}_{n-1,p-1}\exp(s^{\mathrm{gap}\prime\to\mathrm{ali}}))\exp(s^{\mathrm{ali}}_{np} + s^{\mathrm{LA}}_{np}) \\ \alpha^{\mathrm{SA,for,gap}}_{np} = (\alpha^{\mathrm{SA,for,ali}}_{n-1,p}\exp(s^{\mathrm{ali}\to\mathrm{gap}}) + \alpha^{\mathrm{SA,for,gap}}_{n-1,p}\exp(s^{\mathrm{gap}\to\mathrm{gap}}) + \alpha^{\mathrm{SA,for,gap}\prime}_{n-1,p}\exp(s^{\mathrm{gap}\prime\to\mathrm{gap}}))\exp(s^{\mathrm{gap}}_{n}) \\ \alpha^{\mathrm{SA,for,gap}\prime}_{np} = (\alpha^{\mathrm{SA,for,ali}}_{n,p-1}\exp(s^{\mathrm{ali}\to\mathrm{gap}\prime}) + \alpha^{\mathrm{SA,for,gap}}_{n,p-1}\exp(s^{\mathrm{gap}\to\mathrm{gap}\prime}) + \alpha^{\mathrm{SA,for,gap}\prime}_{n,p-1}\exp(s^{\mathrm{gap}\prime\to\mathrm{gap}\prime}))\exp(s^{\mathrm{gap}\prime}_{p}) \end{cases}$$

where $s^{x\to y}$ is the CONTRAlign transition score from the state $x$ to the state $y$, $x,y \in \{\mathrm{ali},\mathrm{gap},\mathrm{gap}\prime\}$, and $s^{\mathrm{gap}}_{n}$ and $s^{\mathrm{gap}\prime}_{p}$ are the CONTRAlign emission scores of the positions $n$ and $p$ aligned with gaps, respectively.

Partition functions $\alpha^{\mathrm{PA},2}_{ijkl}$ and $\alpha^{\mathrm{PA,multi}}_{ijkl}$ recurse into their inner partition functions:

$$\begin{cases} \alpha^{\mathrm{PA},2}_{ijkl} = \exp(s^{\mathrm{PA}}_{ijkl})\sum_{mnop:i<m<n<j,k<o<p<l}\exp(-e^{2}_{ijmn} - e^{2}_{klop})\alpha^{\mathrm{SA,for},(i,j),(k,l),*\to ali}_{m-1,o-1}\alpha^{\mathrm{PA}}_{mnop}\alpha^{\mathrm{SA,back},(i,j),(k,l),ali\to *}_{n+1,p+1} \\ \alpha^{\mathrm{PA,multi}}_{ijkl} = \exp(s^{\mathrm{ali}}_{ik} + s^{\mathrm{ali}}_{jl} + s^{\mathrm{PA}}_{ijkl} - 2e^{\mathrm{CBP}})\alpha^{\mathrm{multi,for},(i,j),(k,l)}_{j-1,l-1} \end{cases}.$$

Here, $\alpha^{\mathrm{SA,for},(i,j),(k,l),*\to ali}_{np} = \alpha^{\mathrm{SA,for},*\to\mathrm{ali}}_{np}$ is the partition function between the pairs $(i+1,n)$ and $(k+1,p)$, whose loops $L_{ij}$ and $L_{kl}$ are 1-loops, and where the state ali is transition to, $\alpha^{\mathrm{SA,back},(i,j),(k,l),ali\to *}_{mo} = \alpha^{\mathrm{SA,back,ali}\to *}_{mo}$ is the partition function between the pairs $(m,j-1)$ and $(o,l-1)$ whose loops $L_{ij}$ and $L_{kl}$ are 1-loops and where the state ali is transition from, and $\alpha^{\mathrm{multi,for},(i,j),(k,l)}_{np} = \alpha^{\mathrm{multi,for}}_{np}$ is the partition function between the pairs $(i+1,n)$ and $(k+1,p)$ whose loops $L_{ij}$ and $L_{kl}$ are multi-loops. A partition function $\alpha^{\mathrm{SA,for},*\to\mathrm{ali}}_{np}$ is calculated as follows:

$$\alpha^{\mathrm{SA,for},*\to\mathrm{ali}}_{np} = \alpha^{\mathrm{SA,for,ali}}_{np}\exp(s^{\mathrm{ali}\to\mathrm{ali}}) + \alpha^{\mathrm{SA,for,gap}}_{np}\exp(s^{\mathrm{gap}\to\mathrm{ali}}) + \alpha^{\mathrm{SA,for,gap}\prime}_{n-1,p-1}\exp(s^{\mathrm{gap}\prime\to\mathrm{ali}})$$

A partition function $\alpha^{\mathrm{multi,for}}_{np}$ consists of three terms: $\alpha^{\mathrm{multi,for}}_{np} = \alpha^{\mathrm{multi,for,ali}}_{np} + \alpha^{\mathrm{multi,for,gap}}_{np} + \alpha^{\mathrm{multi,for,gap}\prime}_{np}$ where $\alpha^{\mathrm{multi,for,ali}}_{np}$, $\alpha^{\mathrm{multi,for,gap}}_{np}$, and $\alpha^{\mathrm{multi,for,gap}\prime}_{np}$ are the partition function $\alpha^{\mathrm{multi,for}}_{np}$ in the cases in which the positions $n$ and $p$ are aligned, the position $n$ is aligned with a gap immediately behind the position $p$, and the position $p$ is aligned with a gap immediately behind the position $n$, respectively. Parition functions $\alpha^{\mathrm{multi,for,ali}}_{np}$, $\alpha^{\mathrm{multi,for,gap}}_{np}$, and $\alpha^{\mathrm{multi,for,gap}\prime}_{np}$ recurse into their forward partition functions:

$$\begin{cases} \alpha^{\mathrm{multi,for,ali}}_{np} = \sum_{mo:m<n,o<p}[(\alpha^{\mathrm{1P,for,ali}}_{m-1,o-1} + \alpha^{\mathrm{multi,for,ali}}_{m-1,o-1})\exp(s^{\mathrm{ali}\to\mathrm{ali}}) + (\alpha^{\mathrm{1P,for,gap}}_{m-1,o-1} + \alpha^{\mathrm{multi,for,gap}}_{m-1,o-1})\exp(s^{\mathrm{gap}\to\mathrm{ali}}) \\ \quad + (\alpha^{\mathrm{1P,for,gap}\prime}_{m-1,o-1} + \alpha^{\mathrm{multi,for,gap}\prime}_{m-1,o-1})\exp(s^{\mathrm{gap}\prime\to\mathrm{ali}})]\exp(-2e^{\mathrm{ABP}})\alpha^{\mathrm{PA}}_{mnop} + (\alpha^{\mathrm{multi,for,ali}}_{n-1,p-1}\exp(s^{\mathrm{ali}\to\mathrm{ali}}) \\ \quad + \alpha^{\mathrm{multi,for,gap}}_{n-1,p-1}\exp(s^{\mathrm{gap}\to\mathrm{ali}}) + \alpha^{\mathrm{multi,for,gap}\prime}_{n-1,p-1}\exp(s^{\mathrm{gap}\prime\to\mathrm{ali}}))\exp(s^{\mathrm{ali}}_{np} + s^{\mathrm{LA}}_{np}) \\ \alpha^{\mathrm{multi,for,gap}}_{np} = (\alpha^{\mathrm{multi,for,ali}}_{n-1,p}\exp(s^{\mathrm{ali}\to\mathrm{gap}}) + \alpha^{\mathrm{multi,for,gap}}_{n-1,p}\exp(s^{\mathrm{gap}\to\mathrm{gap}}) + \alpha^{\mathrm{multi,for,gap}\prime}_{n-1,p}\exp(s^{\mathrm{gap}\prime\to\mathrm{gap}}))\exp(s^{\mathrm{gap}}_{n}) \\ \alpha^{\mathrm{multi,for,gap}\prime}_{np} = (\alpha^{\mathrm{multi,for,ali}}_{n,p-1}\exp(s^{\mathrm{ali}\to\mathrm{gap}\prime}) + \alpha^{\mathrm{multi,for,gap}}_{n,p-1}\exp(s^{\mathrm{gap}\to\mathrm{gap}\prime}) + \alpha^{\mathrm{multi,for,gap}\prime}_{n,p-1}\exp(s^{\mathrm{gap}\prime\to\mathrm{gap}\prime}))\exp(s^{\mathrm{gap}\prime}_{p}) \end{cases}$$

where $\alpha^{\mathrm{1P,for},x}_{np}$ is the partition function $\alpha^{\mathrm{multi,for},x}_{np}$ whose loops $L_{ij}$ and $L_{kl}$ contain one pairing between the pairs $(i+1,n)$ and $(k+1,p)$, respectively. A partition function $\alpha^{\mathrm{1P,for},x}_{np}$ can be computed in a similar manner to the partition function $\alpha^{\mathrm{multi,for},x}_{np}$.

Let $\alpha^{\mathrm{outer}}_{jl}$ be the inside partition function between the pairs $(0,j)$ and $(0,l)$ whose loops $L_{0,N+1}$ and $L_{0,M+1}$ are external. A partition function $\alpha^{\mathrm{outer,for}}_{jl}$ is composed of three terms: $\alpha^{\mathrm{outer,for}}_{jl} = \alpha^{\mathrm{outer,for,ali}}_{jl} + \alpha^{\mathrm{outer,for,gap}}_{jl} + \alpha^{\mathrm{outer,for,gap}\prime}_{jl}$ where $\alpha^{\mathrm{outer,for,ali}}_{jl}$, $\alpha^{\mathrm{outer,for,gap}}_{jl}$, and $\alpha^{\mathrm{outer,for,gap}\prime}_{jl}$ are the partition function $\alpha^{\mathrm{outer,for}}_{jl}$ in the cases in which the positions $j$ and $l$ are aligned, the position $j$ is aligned with a gap immediately behind the position $l$, and the position $l$ is aligned with a gap immediately behind the position $j$, respectively. Partition functions $\alpha^{\mathrm{outer,for,ali}}_{np}$, $\alpha^{\mathrm{outer,for,gap}}_{np}$, and $\alpha^{\mathrm{outer,for,gap}\prime}_{np}$ recurse into their forward partition functions:

$$\begin{cases} \alpha^{\mathrm{outer,for,ali}}_{jl} = \sum_{ik}(\alpha^{\mathrm{outer,for,ali}}_{i-1,k-1}\exp(s^{\mathrm{ali}\to\mathrm{ali}}) + \alpha^{\mathrm{outer,for,gap}}_{i-1,k-1}\exp(s^{\mathrm{gap}\to\mathrm{ali}}) + \alpha^{\mathrm{outer,for,gap}\prime}_{i-1,k-1}\exp(s^{\mathrm{gap}\prime\to\mathrm{ali}}))\alpha^{\mathrm{PA}}_{ijkl} + \\ \quad (\alpha^{\mathrm{outer,for,ali}}_{j-1,l-1}\exp(s^{\mathrm{ali}\to\mathrm{ali}}) + \alpha^{\mathrm{multi,for,gap}}_{j-1,l-1}\exp(s^{\mathrm{gap}\to\mathrm{ali}}) + \alpha^{\mathrm{multi,for,gap}\prime}_{j-1,l-1}\exp(s^{\mathrm{gap}\prime\to\mathrm{ali}}))\exp(s^{\mathrm{ali}}_{jl} + s^{\mathrm{LA}}_{jl}) \\ \alpha^{\mathrm{outer,for,gap}}_{jl} = (\alpha^{\mathrm{outer,for,ali}}_{j-1,l}\exp(s^{\mathrm{ali}\to\mathrm{gap}}) + \alpha^{\mathrm{outer,for,gap}}_{j-1,l}\exp(s^{\mathrm{gap}\to\mathrm{gap}}) + \alpha^{\mathrm{outer,for,gap}\prime}_{j-1,l}\exp(s^{\mathrm{gap}\prime\to\mathrm{gap}}))\exp(s^{\mathrm{gap}}_{j}) \\ \alpha^{\mathrm{outer,for,gap}\prime}_{jl} = (\alpha^{\mathrm{outer,for,ali}}_{j,l-1}\exp(s^{\mathrm{ali}\to\mathrm{gap}\prime}) + \alpha^{\mathrm{outer,for,gap}}_{j,l-1}\exp(s^{\mathrm{gap}\to\mathrm{gap}\prime}) + \alpha^{\mathrm{outer,for,gap}\prime}_{j,l-1}\exp(s^{\mathrm{gap}\prime\to\mathrm{gap}\prime}))\exp(s^{\mathrm{gap}\prime}_{l}) \end{cases}$$

"Backward" inside partition functions $\alpha_{ijkl}^{x,\mathrm{back},y}$ and $\alpha_{uv}^{z,\mathrm{back},w}$ are computed in a similar way to "forward" inside partition functions $\alpha_{ijkl}^{x,\mathrm{for},y}$ and $\alpha_{uv}^{z,\mathrm{for},w}$, for example:

$$\begin{cases} \alpha_{mo}^{\mathrm{SA,back},(i,j),(k,l)} = \alpha_{mo}^{\mathrm{SA,back}} = \alpha_{mo}^{\mathrm{SA,back,ali}} + \alpha_{mo}^{\mathrm{SA,back,gap}} + \alpha_{mo}^{\mathrm{SA,back,gap\prime}} \\ \alpha_{mo}^{\mathrm{SA,back,ali}} = \exp(s_{mo}^{\mathrm{ali}} + s_{mo}^{\mathrm{LA}})(\exp(s^{\mathrm{ali}\to\mathrm{ali}})\alpha_{m+1,o+1}^{\mathrm{SA,back,ali}} + \exp(s^{\mathrm{ali}\to\mathrm{gap}})\alpha_{m+1,o+1}^{\mathrm{SA,back,gap}} + \exp(s^{\mathrm{ali}\to\mathrm{gap\prime}})\alpha_{m+1,o+1}^{\mathrm{SA,back,gap\prime}}) \\ \alpha_{mo}^{\mathrm{SA,back,gap}} = \exp(s_{m}^{\mathrm{gap}})(\exp(s^{\mathrm{gap}\to\mathrm{ali}})\alpha_{m+1,o}^{\mathrm{SA,back,ali}} + \exp(s^{\mathrm{gap}\to\mathrm{gap}})\alpha_{m+1,o}^{\mathrm{SA,back,gap}} + \exp(s^{\mathrm{gap}\to\mathrm{gap\prime}})\alpha_{m+1,o}^{\mathrm{SA,back,gap\prime}}) \\ \alpha_{mo}^{\mathrm{SA,back,gap\prime}} = \exp(s_{o}^{\mathrm{gap\prime}})(\exp(s^{\mathrm{gap\prime}\to\mathrm{ali}})\alpha_{m,o+1}^{\mathrm{SA,back,ali}} + \exp(s^{\mathrm{gap\prime}\to\mathrm{gap}})\alpha_{m,o+1}^{\mathrm{SA,back,gap}} + \exp(s^{\mathrm{gap\prime}\to\mathrm{gap\prime}})\alpha_{m,o+1}^{\mathrm{SA,back,gap\prime}}) \end{cases}.$$

To begin dynamic programming, the initial condition

$$\alpha_{ik}^{\mathrm{SA,for,ali}} = \alpha_{jl}^{\mathrm{SA,back,ali}} = \alpha_{0,0}^{\mathrm{outer,for,ali}} = \alpha_{N+1,M+1}^{\mathrm{outer,back,ali}} = 1$$

is used. (The other inside partition functions $\alpha_{ijkl}^{x}$ and $\alpha_{uv}^{y}$ are set to 0.) The partition function $Z$ is gained from the equivalence $Z = \alpha_{NM}^{\mathrm{outer,for}}$.

#### 1.2 Recursions for efficient computations of outside partition functions $\beta_{ijkl}^{x}$

Let $\beta_{ijkl}^{\mathrm{PA}}$ be the outside partition function between the pair-aligned pairs $(i, j)$ and $(k, l)$. It $\beta_{ijkl}^{\mathrm{PA}}$ consists of three additional components: $\beta_{ijkl}^{\mathrm{PA}} = \beta_{ijkl}^{\mathrm{PA,outer}} + \beta_{ijkl}^{\mathrm{PA,2}} + \beta_{ijkl}^{\mathrm{PA,multi}}$ where $\beta_{ijkl}^{\mathrm{PA,outer}}$, $\beta_{ijkl}^{\mathrm{PA,2}}$, and $\beta_{ijkl}^{\mathrm{PA,multi}}$ are the partition function $\beta_{ijkl}^{\mathrm{PA}}$ whose the pairs $(i, j)$ and $(k, l)$ are in external loops, 2-loops, and multi-loops, respectively. Partition functions $\beta_{ijkl}^{\mathrm{PA,outer}}$, $\beta_{ijkl}^{\mathrm{PA,2}}$, and $\beta_{ijkl}^{\mathrm{PA,multi}}$ can be computed based on partition functions $\alpha_{ijkl}^{x}$ and $\alpha_{uv}^{y}$:

$$\begin{cases} \beta_{ijkl}^{\mathrm{PA,outer}} = \alpha_{i-1,k-1}^{\mathrm{outer,for},*\to\mathrm{ali}}\alpha_{j+1,l+1}^{\mathrm{outer,back,ali}\to*} \\ \beta_{ijkl}^{\mathrm{PA,2}} = \sum_{mnop}\beta_{mnop}^{\mathrm{PA}}\exp(s_{mo}^{\mathrm{ali}} + s_{np}^{\mathrm{ali}} + s_{mnop}^{\mathrm{PA}} - e_{mnij}^{2} - e_{opkl}^{2})\alpha_{i-1,k-1}^{\mathrm{SA,for},*\to\mathrm{ali}}\alpha_{j+1,l+1}^{\mathrm{SA,back,ali}\to*} \\ \beta_{ijkl}^{\mathrm{PA,multi}} = \exp(-2(e^{\mathrm{CBP}} + e^{\mathrm{ABP}}))\sum_{mnop}\beta_{mnop}^{\mathrm{PA}}\exp(s_{mo}^{\mathrm{ali}} + s_{np}^{\mathrm{ali}} + s_{mnop}^{\mathrm{PA}})(\alpha_{i-1,k-1}^{\mathrm{SA,for},*\to\mathrm{ali}}\alpha_{j+1,l+1}^{\mathrm{right,ali}\to*} + \alpha_{i-1,k-1}^{\mathrm{left},*\to\mathrm{ali}}\alpha_{j+1,l+1}^{\mathrm{right,ali}\to*\prime}) \end{cases}$$

where $m < i < j < n$, $o < k < l < p$, and

$$\begin{cases} \alpha_{j+1,l+1}^{\mathrm{right,ali}\to*} = \alpha_{j+1,l+1}^{\mathrm{multi,back,ali}\to*} + \alpha_{j+1,l+1}^{\mathrm{1P,back,ali}\to*} \\ \alpha_{i-1,k-1}^{\mathrm{left},*\to\mathrm{ali}} = \alpha_{i-1,k-1}^{\mathrm{multi,for},*\to\mathrm{ali}} + \alpha_{i-1,k-1}^{\mathrm{1P,for},*\to\mathrm{ali}} \\ \alpha_{j+1,l+1}^{\mathrm{right,ali}\to*\prime} = \alpha_{j+1,l+1}^{\mathrm{right,ali}\to*} + \alpha_{j+1,l+1}^{\mathrm{SA,back,ali}\to*} \end{cases}.$$

Here, partition functions $\alpha_{i-1,k-1}^{x,\mathrm{for},*\to\mathrm{ali}}$ and $\alpha_{j+1,l+1}^{x,\mathrm{back,ali}\to*}$ are the partition functions $\alpha_{i-1,k-1}^{x,\mathrm{for}}$ and $\alpha_{j+1,l+1}^{x,\mathrm{back}}$ considering the transition scores to and from the pairs $(i-1, k-1)$ and $(j+1, l+1)$ under the state ali as the partition function $\alpha_{i-1,k-1}^{\mathrm{SA,for},*\to\mathrm{ali}}$, respectively.

#### 1.3 Efficient computations of accessibilities $p_{ij}^{\lambda^*}$ and $p_{u}^{\lambda}$

Accessibilities $p_{ij}^{\lambda^*}$ can be easily computed while computing probabilities $p_{ijkl}^{\mathrm{PA}}$ because $p_{ij}^{\lambda^*} = \frac{1}{\alpha_{NM}^{\mathrm{outer,for}}}\sum_{kl}\alpha_{ijkl}^{\mathrm{PA}}\beta_{ijkl}^{\mathrm{PA},\lambda^*}$. An accessibility $p_{u}^{1}$ can be computed as

$$p_{u}^{1} = \frac{1}{\alpha_{NM}^{\mathrm{outer,for}}}\sum_{ijklvx:i<u<j,k<v<l,x\in X}\exp(s_{ik}^{\mathrm{ali}} + s_{jl}^{\mathrm{ali}} + s_{ijkl}^{\mathrm{PA}} - e_{ij}^{1} - e_{kl}^{1})\alpha_{uv}^{\mathrm{SA,for},x}\alpha_{u+1,v+1}^{\mathrm{SA,back},x\to*}\beta_{ijkl}^{\mathrm{PA}}$$

where $X = \{\mathrm{ali}, \mathrm{gap}, \mathrm{gap}'\}$. Here, $\alpha_{u+1,v+1}^{\mathrm{SA,back},x\to*}$ is the partition function $\alpha_{u+1,v+1}^{\mathrm{SA,back}}$ considering the transition scores from the positions $u+1$ and $v+1$ under the state $x$ as the partition function $\alpha_{u+1,v+1}^{\mathrm{SA,back,ali}\to*}$. An accessibility $p_{u}^{2}$ can be computed as

$$p_{u}^{2} = (\sum_{mnij:m<u<i<j<n} p_{mnij}^{2} + \sum_{mnij:m<i<j<u<n} p_{mnij}^{2})$$

where $p_{mnij}^{2}$ is the probability that the pairing positions $i$ and $j$ are in the loop $L_{mn}$, which constitutes the accessibility $p_{ij}^{2}$. An accessibility $p_{u}^{\mathrm{multi}}$ can be computed as

$$p_{u}^{\mathrm{multi}} = \frac{\exp(-2e^{\mathrm{CBP}})}{\alpha_{NM}^{\mathrm{outer,for}}}(\sum_{ijklvx:i<u<j,k<v<l,x\in X}\exp(s_{ik}^{\mathrm{ali}} + s_{jl}^{\mathrm{ali}} + s_{ijkl}^{\mathrm{PA}})(\alpha_{uv}^{\mathrm{SA,for},x}\alpha_{u+1,v+1}^{\mathrm{multi,back},x\to*} + \alpha_{uv}^{\mathrm{1P,for},x}\alpha_{u+1,v+1}^{\mathrm{right},x\to*} + \alpha_{uv}^{\mathrm{multi,for},x}\alpha_{u+1,v+1}^{\mathrm{right},x\to*\prime})\beta_{ijkl}^{\mathrm{PA}}.$$

Here, $\alpha_{u+1,v+1}^{\mathrm{right},x\to*}$ and $\alpha_{u+1,v+1}^{\mathrm{right},x\to*\prime}$ are the partition functions $\alpha_{u+1,v+1}^{\mathrm{right}} = \alpha_{u+1,v+1}^{\mathrm{multi,back}} + \alpha_{u+1,v+1}^{\mathrm{1P,back}}$ and $\alpha_{u+1,v+1}^{\mathrm{right}\prime} = \alpha_{u+1,v+1}^{\mathrm{right}} + \alpha_{u+1,v+1}^{\mathrm{SA,back}}$ considering the transition scores from the positions $u+1$ and $v+1$ under the state $x$ as the partition function $\alpha_{u+1,v+1}^{\mathrm{SA,for},*\to\mathrm{ali}}$. An accessibility $p_{u}^{\mathrm{outer}}$ can be computed as

$$p_{u}^{\mathrm{outer}} = \frac{1}{\alpha_{NM}^{\mathrm{outer,for}}}\sum_{vx:x\in X}\alpha_{uv}^{\mathrm{outer,for},x}\alpha_{u+1,v+1}^{\mathrm{outer,back},x\to*}.$$
